# Combinations of DIPs and Dprs control organization of olfactory receptor neuron terminals in Drosophila

**DOI:** 10.1101/316109

**Authors:** Scott Barish, Sarah Nuss, Ilya Strunilin, Suyang Bao, Sayan Mukherjee, Corbin Jones, Pelin C. Volkan

## Abstract

In *Drosophila*, 50 classes of olfactory receptor neurons (ORNs) connect to 50 class-specific and uniquely positioned glomeruli in the antennal lobe. Despite the identification of cell surface receptors regulating axon guidance, how ORN axons sort to form 50 stereotypical glomeruli remains unclear. Here we show that the heterophilic cell adhesion proteins, DIPs and Dprs, are expressed in ORNs during glomerular formation. Each ORN class expresses a unique combination of *DIPs*/*dprs*, with neurons of the same class expressing interacting partners, suggesting a role in class-specific self-adhesion ORN axons. Analysis of DIP/Dpr expression revealed that ORNS that target neighboring glomeruli have different combinations, and ORNs with very similar DIP/Dpr combinations can project to distant glomeruli in the antennal lobe. Perturbations of *DIP*/*dpr* gene function result in local projection defects of ORN axons and glomerular positioning, without altering correct matching of ORNs with their target neurons. Our results suggest that context-dependent differential adhesion through DIP/Dpr combinations regulate self-adhesion and sort ORN axons into uniquely positioned glomeruli.

## Introduction

One of the most complex biological systems in nature is the human brain, which contains an estimated 86 billion neurons wired to make approximately 100 trillion synaptic connections (1). Our ability to experience the world, solve complex problems, create, and behave, all depends on the molecular, morphological, and functional diversity among the billions of neurons, and their wiring patterns established during development. Once neurons are born, they extend their axons long distances until they reach a target site where neurons in different circuits sort into specific structures and select target neurons or muscle cells with which to make synaptic connections (2,3). The expression of genes, particularly those encoding cell surface receptors (CSRs), relay attractive or repulsive cues to regulate each step of this wiring program. Mutations affecting these programs are associated with numerous neurodevelopmental and neuropsychiatric disorders, as well as many known brain cancers (4–7). It is currently thought that CSRs and their ligands act in combinations, as well as different concentration gradients to regulate different steps of circuit assembly during development (2). While individual examples of CSRs directing axon guidance and connectivity are well known and evolutionarily conserved, how they act in combinations to coordinate large scale organizational patterns among a diverse set of neurons within a circuit remains poorly understood.

The *Drosophila* olfactory system provides an excellent system to identify these determinants, where neurons belonging to 50 different olfactory receptor neuron (ORN) classes sort out and synapse with their target projection neurons within 50 unique glomeruli in the antennal lobe (8). Each ORN class is defined by the exclusive expression of typically a single olfactory receptor gene from approximately 80 possibilities in the genome (9,10). ORNs of the same class converge their axons into a distinctly positioned and class-specific glomerulus in the antennal lobe, generating 50 unique glomeruli targeted by the 50 different classes of ORNs (11–13). The molecular parameters that establish glomerular organization are not known.

The organizational logic of the peripheral olfactory system is conserved in many species including mammals. For example, mice have over a million ORNs grouped into ∼1000 classes based on the expression of a single OR gene from over 1000 possibilities in the genome (14). ORNs of the same class converge their axons onto the same glomerulus in the olfactory bulb, where they synapse with 2^nd^ order mitral/tufted cell dendrites (15). Mammalian olfactory receptors, through ligand-dependent and - independent G-protein coupled signaling, were shown to differentially regulate the expression of CSRs to direct glomerular positioning of ORNs (16,17). *Drosophila* olfactory receptors however, are ligand gated cation channels and do not contribute to glomerular organization (18). Thus, how the *Drosophila* olfactory system coordinately positions 50 classes of ORNs into 50 distinct glomeruli requires further study. With its diverse yet workable amount of ORN classes, the availability of the wiring map, and reporters for all ORNs, the *Drosophila* olfactory system is a powerful model to gain a systems level understanding of how 50 ORN classes can coordinate highly stereotyped organizational patterns in the brain.

Adult ORNs in Drosophila are born from the asymmetric division of precursors located in the larval antennal disc, which, during pupal metamorphosis, becomes the adult antenna (19,20). ORN axons reach the future antennal lobe by 16-18 hours after puparium formation (APF) (21,22). No glomeruli can be seen at these stages (22). Most glomeruli are clearly detectable by antibodies against N-Cadherin, around 40 hours APF, and fully separate into distinct structures by 48-50 hrs APF (22). Several genes have been shown to regulate each step of ORN axon guidance, such as Semaphorins/Plexins (axonal tract selection, interclass repulsion, (23,24)), Dscam and Robos (axon targeting, (25,26)), N-Cadherin (intraclass attraction, (27,28)), and Teneurins and Tolls (ORN-PN matching, (29–31)). The vast majority of these proteins however, work very broadly and are required for the proper targeting of most or all ORN classes (20,32). It is therefore still unclear how axons of 50 different ORN classes organize themselves to form 50 uniquely positioned and structured glomeruli. This is likely due to the complex and combinatorial nature of the molecular interactions underlying ORN wiring patterns, which includes programs for axon-axon sorting/positioning and target specificity in the antennal lobe.

Here we identify the Defective proboscis response (Dpr) family proteins and their heterophilic binding partners Dpr Interacting Proteins (DIPs) as novel regulators of glomerular positioning and structure in the *Drosophila* olfactory system. Each ORN class expresses a unique combination of DIP/Dprs starting at stages dedicated to glomerular formation. Interestingly, interacting DIP/Dpr partners are generally, but not exclusively, found in the same ORN class, likely aiding self-adhesion among axons of the same ORN class, while also sorting from others. Mathematical analysis of class-specific DIP/Dpr expression showed that ORNs with very similar DIP/Dpr combinations, can end up in distant glomeruli in the antennal lobe, and ORNs targeting neighboring glomeruli can have very different combinations. DIPs/Dprs control the class-specific positioning of ORN axon terminals and their glomerular morphology. Perturbations to DIP/Dpr combinations are associated with context-dependent, and local disruptions of glomerular morphology and positioning and in some cases, invasions of neighboring glomeruli, without changing ORN-PN matching. These results suggest that differential adhesion among local ORN axons, mediated by DIPs/Dprs, determines the position and morphology of each glomerulus. Our results demonstrate how combinatorial action of many interacting CSRs can generate differential adhesive forces as a strategy to coordinately organize axons of all circuits within a neural system.

## Results

To identify candidate genes that may be involved in the establishment of 50 class specific glomeruli, we analyzed our previously reported antennal transcriptome data from four stages of development: 3^rd^ instar larval antennal discs (3L), 8 hrs after puparium formation (APF) antennal discs (p8), 40 hrs APF antennae (p40) and adult antennae (Adult) (33,34). Axon guidance, glomerular sorting, and ORN-PN matching are inherently temporal processes that occur in a specific developmental order (8). We therefore mined this dataset to identify novel regulators of wiring specificity of ORNs, whose expression overlapped with the timing of glomerular formation. We queried ∼250 Flybase annotated cell surface receptors (CSRs) and analyzed their developmental expression patterns using hierarchical clustering to group genes based upon their developmental expression patterns (Fig 1A, Table S1). We found 8 clusters of CSRs with distinct expression patterns (Fig 1A, B). Two broad patterns emerged from this analysis, genes that are expressed at constant levels throughout (Clusters 3, 5, 6, and 8, Fig 1A, B), such as Sema-1a/b and Dscam1, and genes that are weakly expressed early in development and increase in later stages (Cluster 1 and 2, Fig 1A, B), such as Robo3 (Table S1). Cluster 7 contained many genes that were highly expressed at the first three stages of development but decreased at the adult stage, and Cluster 4 contained genes whose expression peaked at 40 hrs APF (p40). Known regulators of ORN wiring grouped into Clusters 7 and 8 as well as Cluster 1, meaning that they were expressed highly throughout development or lowly expressed early and increased their expression at the later stages (Fig 1A-C) consistent with their roles in ORN axon guidance, which begins very early in olfactory system development.

**Figure 1:**
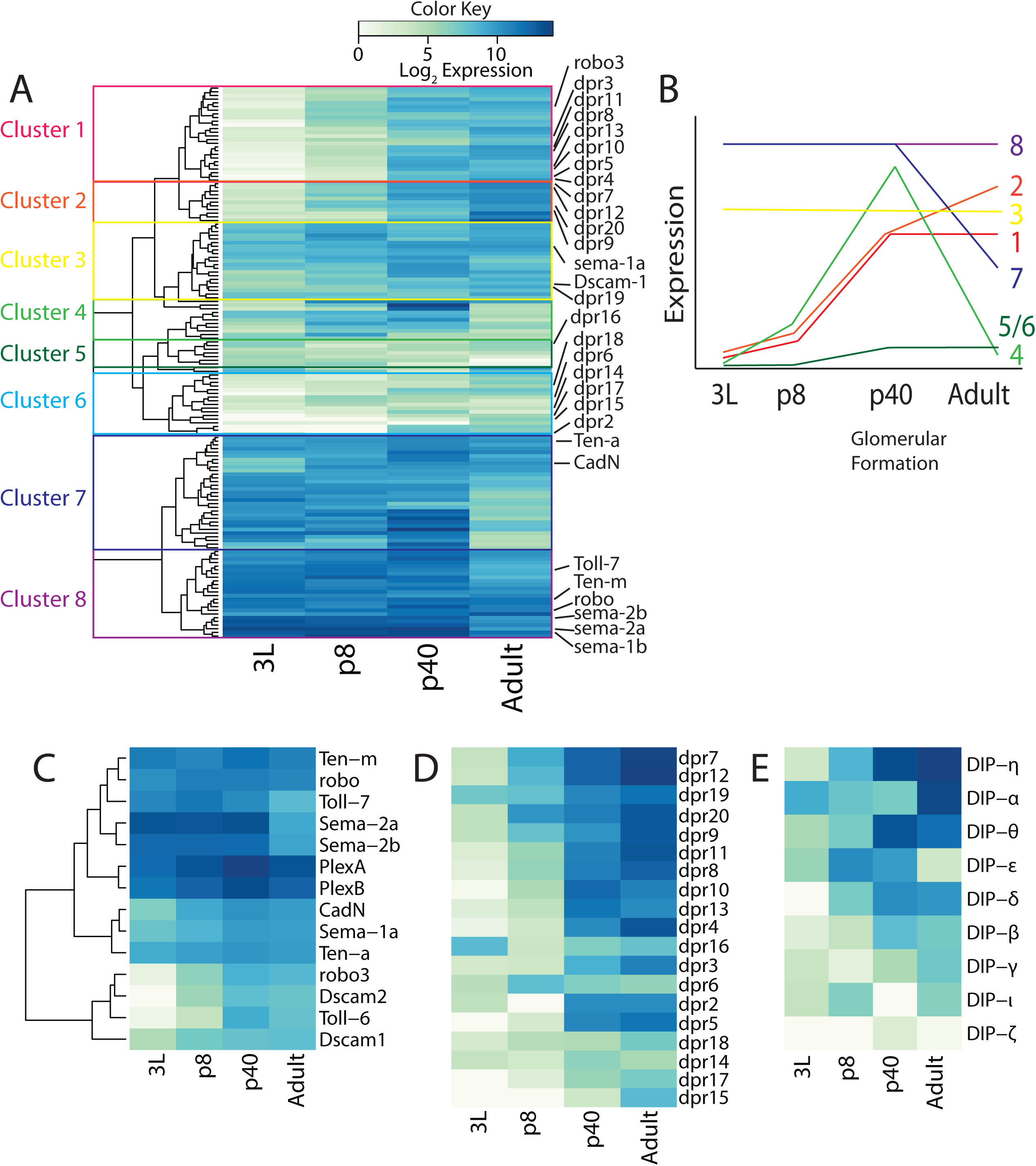
Transcriptional profiles of CSR genes in the developing *Drosophila* olfactory system. Heatmaps representing normalized log_2_ expression values for CSRs. Highly expressed genes are represented in darker colors, and genes with lower expression levels are represented with lighter colors. A) Hierarchical clustering of the developmental expression patterns of in the olfactory system groups genes into clusters. Known regulators of ORN wiring and members of the *dpr* family are highlighted. B) Schematic showing expression profile of genes in each cluster at the 4 developmental stages. C) Hierarchical clustering of the developmental expression patterns of known regulators of ORN wiring reveals two major expression patterns: high expression at all stages, and low expression at early stages followed by high expression at later stages. Expression patterns of *dpr* (D) and *DIP* (E) genes ordered from highest expression across all stages to lowest. Most *DIP/dpr* genes are expressed primarily at later developmental stages (p40 and Adult).

### DIP and Dprs are expressed in late stages of ORN wiring

Most ORN axons surround both antennal lobes, begin and complete glomerular formation by 30 hrs, 40 hrs, and 48 hrs APF, respectively (32,34). We therefore hypothesized that genes that were highly expressed at the two later stages of development (p40 and Adult) particularly p40, but lowly expressed at the two early stages (3L and p8) were more likely to be involved in class-specific glomerular formation. Our analysis of CSR expression profiles identified that the majority of Dpr family of CSR proteins and their binding partner DIP proteins were expressed at p40 and in adult antennae but showed low or no expression during earlier stages of antennal development (Fig 1D, E). These results pointed to a possible role for DIP/Dpr family members in glomerular formation.

### DIPs and Dprs are expressed in a combinatorial code in ORNs

Dprs and their binding partners DIPs are members of the Ig superfamily of proteins and each contain 2-3 Ig extracellular domains (35). Members of the Ig superfamily of proteins, such as Dscam, and Kirrels, are well established to control axon guidance and sorting in other systems (17,36). DIPs/Dprs themselves have recently been shown to direct synaptic target matching in the *Drosophila* eye (35,37). In addition, the relatively large number of *DIP/dpr* genes (9 and 21 respectively) make them good candidates to contribute to a combinatorial code of CSRs that direct wiring specificity for each class of ORNs.

Although our RNA-seq data establish temporal patterns of *DIP*/*dpr* expression in the developing olfactory system, it does not inform us of the ORN class-specific expression of each *DIP*/*dpr* gene. To investigate which ORN classes express each *DIP*/*dpr*, we used MIMIC insertion derived GAL4 lines for each *DIP* and *dpr* to drive UAS>STOP>GFP with *eyeless* driven *flippase* to express GFP specifically in ORNs (35,37,38). Because the glomerular positions for all ORN classes have been mapped, we were able to determine the ORN classes that express each *DIP/dpr* gene, based on GFP expression in the antennal lobe (Fig 2, S1A-L). These analyses showed that each ORN class has a unique *DIP/dpr* expression profile (Fig 3A). Some *DIPs/dprs* are expressed very broadly across ORN classes (*DIP-η*, Fig 2H), and others that are expressed in fewer ORN classes (*DIP-β* Fig 2E). These results suggest that the unique combination of DIPs and Dprs in each ORN class might direct ORN class specific glomerular formation.

**Figure 2:**
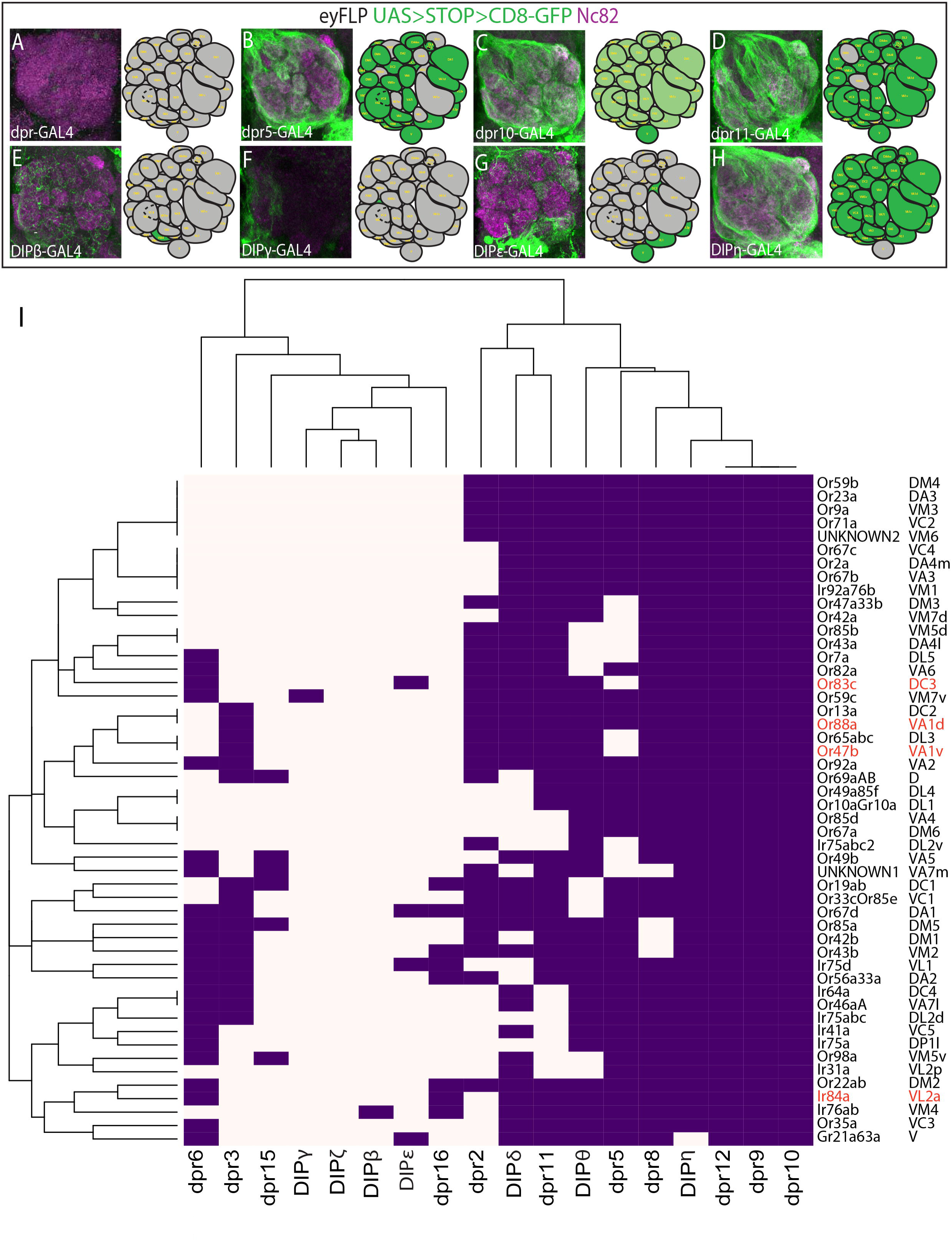
Combinatorial and ORN class specific expression patterns of *DIP* and *dpr* genes. *DIP* and *dpr*-*GAL4s* were used to drive *UAS>STOP>GFP* in ORNs with *eyFLP* (green, A-H) to visualize ORN axons with staining for the neuropil (magenta). Each gene is expressed in a unique set of glomeruli and ORN classes. A map of the expression pattern of gene in the antennal lobe is provided in the right panel (A-H). Results for all genes are summarized in Supplemental Figure 1. (I) Hierarchical bi-clustering of DIP/Dpr expression patterns by ORN class reveals that each class of neurons expresses a unique combination of DIPs/Dprs. ORN classes highlighted in red neighbor each other in the antennal lobe and are further analyzed in subsequent figures.

**Figure 3:**
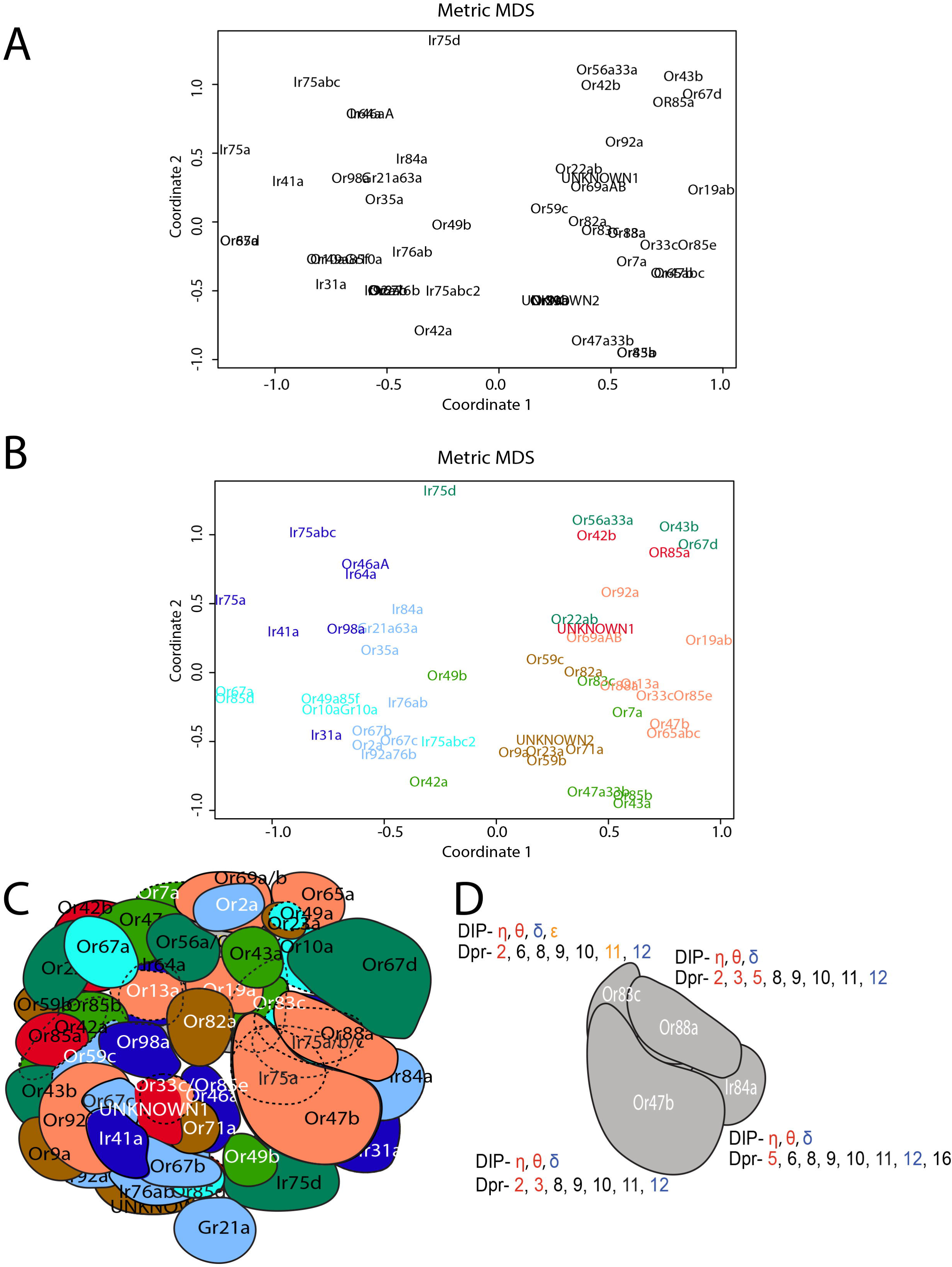
Multidimensional scaling clusters ORN classes by DIP/Dpr expression pattern, which groups classes targeting distant glomeruli. (A) Multidimensional scaling analysis clusters ORN classes based upon DIP/Dpr expression and glomerular distances. (B) K-means clustering was used to determine which ORN classes shared the most similar DIP/Dpr expression profiles in the MDS analysis. Classes that were clustered together were assigned the same color. (C) ORN classes clustered in B tend to target distant glomeruli. Coloring of glomeruli matches clusters in B. (D) Schematic of four glomeruli (Or47b, Or88a, Or83c, and Ir84a) that will be analyzed in subsequent figures. The expression code for each glomerulus is highlighted with each receptor-ligand pair displayed in matching colors.

Next, we analyzed expression of *DIPs/dprs* in the antenna to confirm the expression patterns we observed in the antennal lobe corresponded to expression in ORN cell bodies. To do this we used *DIP/dpr-GAL4*s to drive the expression of *UAS-CD8GFP* (Fig S1M-W). Consistent with our RNA-seq analysis, most of the genes we analyzed showed expression in ORNs consistent with glomerular expression patterns (Fig S1M-W). A small number of genes did not show expression in the antenna (Fig S1M-W), which suggests that they are instead expressed in local interneurons. We conclude that the majority of *DIPs/dprs* are robustly expressed in ORNs but a few are expressed in local interneurons.

Glomeruli in the antennal lobe finalize their positioning and morphology between 40-50 hrs APF (22). After this time, glomeruli enlarge prior to eclosion, but do not otherwise alter their shape (22). Thus, we predicted that the expression of *DIPs/dprs* at this time would reveal how they contribute to shaping glomerular morphology and positioning. We analyzed the expression pattern of some *DIPs/dprs* specifically in ORNs and found that many were expressed in a developmentally dynamic manner. For example, *DIP-η*, which is expressed in all ORNs except Gr21a neurons in the adult, is missing from some classes at 50 hours APF (Fig S2). In addition, *dprs 9, 10,* and *11* were expressed in a smaller number of classes as compared to their adult expression patterns (Fig S2E-G). Other *DIPs/dprs* displayed similar or identical expression patterns to their adult expression, particularly those that had sparse expression patterns at both stages, such as *dpr16* and *DIP-γ* (Fig S2I, K). Together, our results suggest that *DIPs* and *dprs* are expressed at the final stages of glomerular formation and may play multiple roles in glomerular positioning and formation based upon their dynamic expression patterns.

Previous studies have shown that the expression of *dprs* in photoreceptor cells and their interacting DIPs in the target lamina neurons contribute to synaptic partner matching in the *Drosophila* visual system (35,37). In order to ask whether DIPs/Dprs play a similar role in synaptic partner matching between ORNs and their target PNs we next analyzed the expression patterns of each *DIP/dpr* in PNs (Fig S3). We drove *UAS-DenMark-RFP* using *DIP/dpr-GAL4s*, which specifically labels postsynaptic dendrites (39), thereby labeling postsynaptic processes of PNs in the antennal lobe. Similar to ORNs and in agreement with recent single cell RNAseq studies, some of the genes were expressed in specific subsets of PNs (*DIP-η* and *DIP-γ,* Fig S3), while others were expressed much more broadly (*dpr10*, Fig S3E) (40). Several however, were not expressed in PNs at all (*dpr12*, Fig S3F). Unlike in previous reports, we did not observe any obvious pattern of DIPs/Dprs indicative of pre- and post-synaptic target matching between ORNs and their corresponding PNs. Taken together, our expression data suggest that DIPs/Dprs may be involved in ORN-ORN axon interactions.

### Mathematical analysis based on DIP/Dpr profiles of ORNs cluster ORN classes that target distant glomeruli

Analysis of DIP/Dpr patterns in each ORN class, showed that interacting DIP-Dpr pairs are expressed in the same ORN classes (Fig 2I, 3D). DIPs and Dprs heterophilically interact in a combinatorial code that has been previously described (Fig S4) (37). We also found some ORN classes additionally express *dpr* ligands without their *DIP* receptors, which might regulate interactions with DIP receptors on neighboring ORN axons during development (Fig 2I, 3D, see discussion). To determine which ORN classes are most similar based upon their DIP/Dpr gene signatures from the GAL4 expression patterns we applied hierarchical biclustering to the data (Fig 2I). This analysis clustered specific classes of ORNs together (Fig 2I). Typically, ORN classes that target neighboring glomeruli did not cluster together in this analysis. An exception to this are Or47b, Or65a, and Or88a ORNs, which reside in the same sensillum and target neighboring glomeruli. Instead, ORNs targeting distant glomeruli expressed similar DIP-Dpr profiles. (Fig 2I).

Given our results, we predicted that class-specific axon sorting may be dictated by DIP/Dpr combinations and might regulate glomerular positioning and formation. To test whether the unique DIP/Dpr relationships among ORN classes can describe their relative glomerular positions in the antennal lobe, we performed multidimensional scaling (MDS) on the expression pattern data set (Fig 3A). We used the expression of *DIP/dpr* genes in each ORN class as the input variables. The genes and gene combinations defining each ORN class were used to create a matrix of “distances” between ORN classes. We predicted that this statistical analysis would group similar combinations of DIP/Dpr profiles for each ORN class, and sort them to distinct coordinates with respect to one another. Once the MDS results were plotted, we used k-means clustering to determine clustered ORN classes, and color coded them on the MDS plot (Fig 3B). We next color coded antennal lobe glomeruli based upon the clusters derived from the k-means analysis (Fig 3C). Plotting ORN classes using this method revealed that ORNs with very similar DIP/Dpr combinations, tended to target distant glomeruli within the antennal lobe. In addition, neighboring glomeruli have different combinations of DIP/Dprs. These results suggest that these differences can drive ORN class-specific adhesion, consequently sorting them from the axon terminals of other ORN classes, which themselves have to self-adhere (Fig 3B, C).

The DIP-Dpr profiles of each ORN might arise as a result of developmental programs assigning each ORN its sensilla type and subtype identity, as well as a result of Notch signaling that assigns sensory and wiring identities (19,41). Thus, we re-examined the MDS analysis to visualize how sensilla types and Notch state are represented on the MDS plot. We first examined sensilla types, basiconics (large, thin, small, and palp), trichoids, coeloconics, and intermediates (Fig S5A). When we removed basiconics from the plot however, we observed that DIP/Dpr profiles of coeloconic and trichoid sensilla occupied unique positions on the MDS plot (Fig S5B), suggesting DIP/Dpr combinations of ORNs within either sensilla type are more similar among the ORNs of the same sensilla type compared to the others (Fig 3B). This pattern is consistent with the observation that trichoid ORN classes target more dorsal and anterior regions of the antennal lobe, whereas coeloconic classes target more ventral and medial regions (42). Basiconic sensilla ORNs did not show such clear segregation patterns in this MDS plot (Fig S5C) (42). These results suggest that DIP/Dpr profiles can be regulated by programs of sensilla type identity for trichoid and coeloconic ORNs, but not for basiconic ORNs.

Next, we analyzed whether the Notch states segregate DIP/Dpr profiles of ORN classes in the MDS analysis. Within each sensilla, sensory and glomerular targeting fates of ORN pairs are segregated using Notch signaling, where each fate is associated with a coordinated Notch ON or Notch OFF state (19,43). Labeling classes as Notch-ON or Notch-OFF on the plot did not reveal any noticeable patterns within sensilla types (Fig S5D, E). These data suggest that the expression of *DIPs/dprs* are likely not under the control of Notch-Delta signaling.

Even though this plot reveals that ORNs with different DIP/Dpr combinations project to neighboring glomeruli, it is not complete, and contains overlaps with some ORNs with identical DIP/Dpr profiles. Further refinement of the plot will require identification and analysis of the entire CSR profiles of ORN classes, their modes of interaction, and function. Regardless, our analysis highlights the relative relationships among particular classes of ORNs based upon their *DIP*/*dpr* expression profiles, and supports the hypothesis that DIPs/Dprs can regulate class-specific sorting of ORN axons into distinctly positioned glomeruli.

### Perturbations to *DIP/dpr* code are associated with local ORN axon terminal positioning defects

Based on the class specific expression patterns of *DIPs/dprs*, we suspected that knocking down individual genes would cause disruptions to class-specific sorting of ORN axons and glomerular formation. To investigate whether DIPs/Dprs are required for appropriate ORN axon projections in the antennal lobe, we used RNAi-mediated knock down against a panel of *DIPs/dprs* using the *peb-GAL4*, which is expressed in all ORNs beginning early in olfactory system development. Analysis Or47a, Or47b, and Gr21a axon terminals in the antennal lobe did not reveal any significant changes in glomerular projection patterns (Fig S6). We predicted that this is likely due to the combinatorial, redundant, and context-dependent function of the DIP/Dpr proteins expressed in specific ORNs or their glomerular neighbors.

To simplify our analysis and better understand the combinatorial function of DIPs/Dprs we chose to focus our experiments on a group of four neighboring glomeruli, VA1v (Or47b), VA1d (Or88a), VL2a (Ir84a), and DC3 (Or83c). We initially focused on these neighboring glomeruli because DIPs/Dprs are membrane bound proteins with short-range, heterophilic interactions (35,37). Among these four classes, Or47b and Or88a ORNs have very similar DIP/Dpr profiles that also clustered in the MDS plots, and only differ in a few Dprs (Fig 4B-D). Thus, we reasoned that ORN projections into these two glomeruli may be particularly sensitive to genetic manipulation. For example, Or47b ORNs are positive for *DIP-η, DIP-θ, DIP-δ, dpr2, dpr3, dpr8, dpr9, dpr10, dpr11,* and *dpr12*, whereas Or88a ORNs express the same genes plus *dpr5* (Fig 2I, 3D). Ir84a and Or83c ORNs on the other hand express *DIP-η, DIP-θ, DIP-δ, dpr5, dpr6, dpr8, dpr9, dpr10, dpr11, dpr12* and *dpr16*; and *DIP-η, DIP-θ, DIP-δ, DIP-ε, dpr3, dpr6, dpr8, dpr9, dpr10, dpr11,* and *dpr12,* respectively (Fig 2I, 3D). This code differentiates each class of ORN axons from each other and provides candidate genes to manipulate singly or in combination to investigate the role of DIPs/Dprs in controlling ORN axon projection patterns in the antennal lobes.

**Figure 4:**
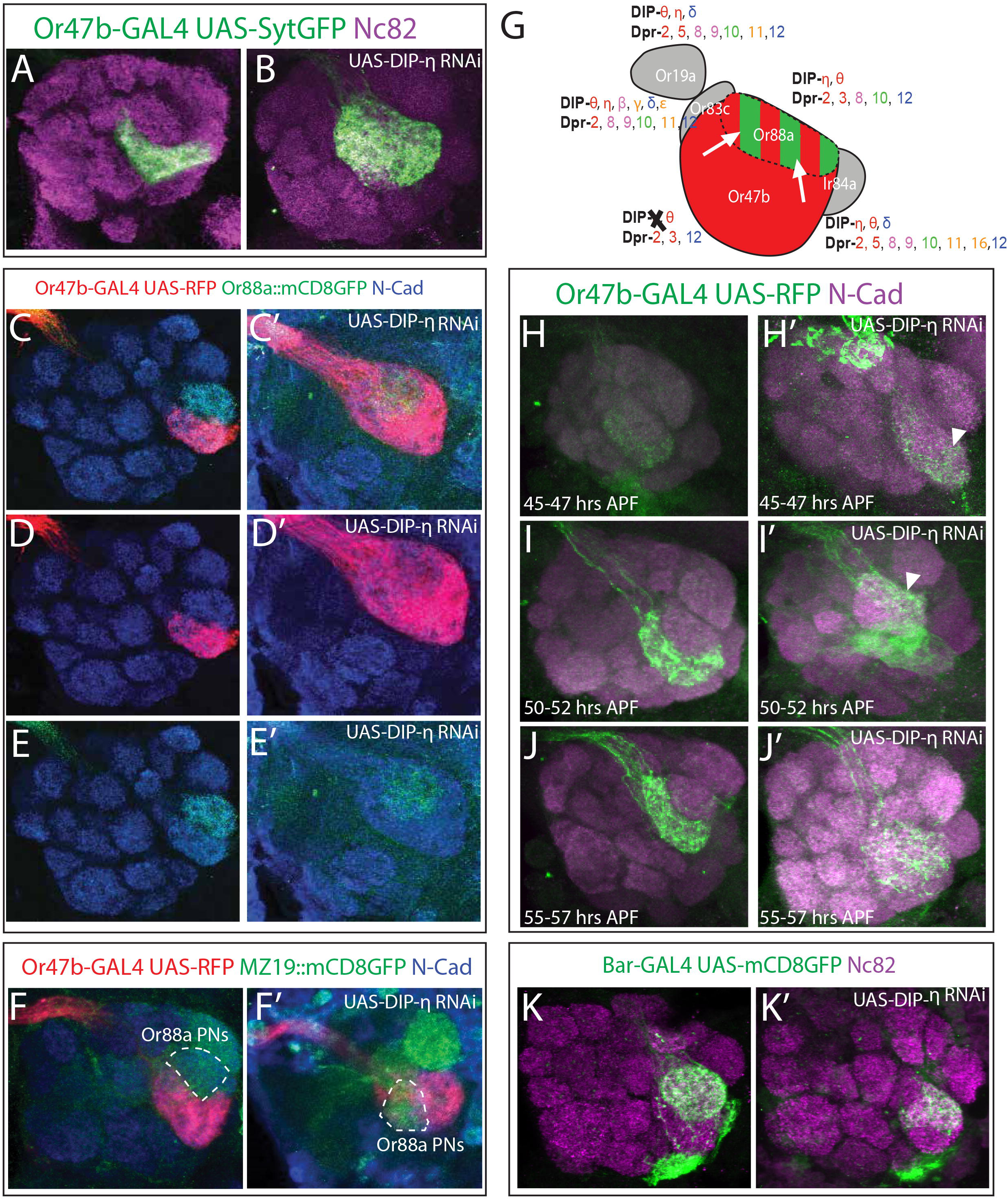
Cell-autonomous knock down of DIP-η in Or47b ORNs disrupts sorting of Or47b axons from Or88a glomerulus. (A, B) Knock down of *DIP-η* with the *Or47b-GAL4* driver, also used to drive *UAS-synaptotagmin-GFP* (green), causes Or47b axons to expand and invade a neighboring glomerulus. (C-E’) Simultaneous labeling of Or47b (red) and Or88a (green) axons reveals that knock down of *DIP-η* causes Or47b ORN axons to invade the Or88a ORN target glomerulus, while Or88a ORN axons retain their ability to coalesce into a glomerulus despite intermingling with Or47b axons. (F-F’) Co-labeling of Or47b axons (red) and MZ19 expressing PNs (green). During knock down of *DIP-η,* MZ19 expressing PNs do not invade the VA1v glomerulus. (G) Schematic of axon sorting phenotypes in *DIP-η* knock down. When *DIP-η* is specifically ablated from Or47b ORNs (black X), Or47b ORN axons (red) invade the Or88a ORN target glomerulus (white arrows) and intermingle with Or88a ORN axons (red/green striping). (H-J’) Developmental analysis of *DIP-η* RNAi phenotype with Or47b axons labeled in green and N-Cadherin in magenta, at 45-47 hrs APF (H-H’), 50-52 hrs APF (I-I’), and 55-57 hrs APF (J-J’). Disruptions to VA1v glomerular morphology (arrowheads) can be detected as early as 45-47 hrs APF (H’) and larger expansions, as seen in the adult, can be observed by 55-57 hrs APF (J’). (K-K’) Knock down of *DIP-η* using the *Bar-GAL4,* which expresses in Or88a, Ir84a, and Ir75d ORNs (33). Anterior sections of the antennal lobe are shown, and disruption to the VA1d glomerulus can be observed (K’).

Interestingly, each ORN class, generally expresses interacting DIP/Dpr partners, suggesting that not only can DIP/Dprs contribute to sorting glomeruli, but also heterophilic interactions among ORN axons of the same class can lead to self-adhesion. We hypothesized that manipulating the DIP/Dpr code in these specific ORN classes would cause local projection defects, such as expansion and splits in glomeruli, as wells as defects in glomerular positioning, within this cluster. To test this hypothesis, we drove RNAi against single DIPs/Dprs in specific classes or against combinations in all ORNs. *DIP-η* is expressed in all four ORN classes projecting into these glomeruli, and its knock down in all ORNs does not result in any defects (Fig S6D). Based upon the specific expression of *dpr3,* a *DIP-η* interaction partner, at 40-50 hrs APF (Fig S2A), we reasoned that perturbation of *DIP-η* in these ORNs specifically would cause glomerular defects. Thus, we began by knocking down *DIP-η* in only Or47b neurons using the *Or47b-GAL4*. We found that knockdown of *DIP-η* specifically disrupted the positioning and morphology of the VA1v glomerulus causing it to dramatically expand dorsally (Fig 4A, B). This phenotype was extremely penetrant, with 85% of individuals displaying expansion of the VA1v in one or both antennal lobes, (n=12) but wildtype individuals never displayed any invasions (n=10, Fig S7C). We observed that the expansion of the Or47b glomerulus was not random, and always appeared to overtake the VA1d glomerulus, which is dorsally adjacent to the VA1v glomerulus (12). Co-labeling of Or47b and Or88a ORN axons revealed that Or47b ORN axons indeed expand towards the VA1d glomerulus, but this does not disrupt the ability of Or88a ORN axons to form a glomerulus (Fig 4C-E’). Instead, the Or88a axons occupy one part of the now enlarged VA1v glomerulus (Fig 4C’-E’).

Previous reports of Or47b axon invasion of the VA1d glomerulus has been caused by disruptions to ORN specification programs within at4 sensillum (44). To test whether ORN specification is altered in *DIP-η* knock down, we labeled Or47b and Or88a ORN cell bodies with RFP and GFP, respectively (Fig S7D, E). Comparison of Or47b ORN numbers in the antennae showed no significant difference between wildtype and *DIP-η* knock down antennae (56.7 vs 57.8, p=0.64, Fig S7F), suggesting that the expansion of VA1v glomerulus in *DIP-η* knock down flies is not due to an increase Or47b ORNs. We also counted the number of Or88a ORNs in Or47b specific *DIP-η* RNAi knock down (Fig S7D, E). We found a significant reduction in the number of Or88a neurons (46.8 vs 18.2, p<0.001, Fig S7F). This result was unexpected as we drove RNAi against *DIP-η* in Or47b neurons and RNAi expression begins well after the birth of both Or47b and Or88a neurons. This effect is unlikely to alter wiring patterns however, as it has been previously reported that ablation of ORNs does not affect the connectivity of ORNs that target neighboring glomeruli (45). We therefore conclude that *DIP-η* is required cell-autonomously, after the onset of olfactory receptor expression, to organize Or47b ORN axon projections and position them with respect to Or88a ORN axons to form two distinct glomeruli. *DIP-η* may also non-autonomously affect the survival of Or88a neurons.

Because the expansion of Or47b ORN axon terminals towards the VA1d glomerulus could be due to a defect in ORN-PN matching, we next investigated the behavior of VA1d PNs using the *MZ19* reporter that labels DA1, DC3, and VA1d PNs in Or47b specific knock down of *DIP-η* (Fig 4F, F’). We expected that a defect in ORN-PN matching would cause VA1d or DC3 PNs to invade reciprocally into the VA1v glomerulus. Co-labeling of Or47b axons and *MZ19* PNs shows that while VA1d PNs are occasionally mis-located relative to the VA1v glomerulus, they maintain glomerular integrity and do not invade into the VA1v glomerulus (n=14, Fig 4F’). Thus, we conclude that loss of DIP-η function does not lead to ectopic matching in Or47b ORNs with VA1d PNs. These results suggest that the projection defects in Or47b specific *DIP-η* knock down are likely due to disrupted axon-axon interactions between Or47b and Or88a (and possibly Or83c and Ir84a) ORNs, but not ORN-PN matching.

To investigate when the *DIP-η* knock down phenotype arises, we conducted a time series analysis beginning at 45 hrs APF just after the onset of Or47b expression in the antenna (34). At 45 and 50 hrs APF, we observed that the VA1v glomerulus failed to properly orient in relation to the VA1d glomerulus (n=5 80% penetrant and n=7 57% penetrant respectively Fig 4H-I’, Fig S6G). This included splitting of the VA1v glomerulus. By 55 hrs APF however, we observed a more complete expansion of the VA1v glomerulus to overtake the VA1d glomerulus (n=12 75% penetrance, Fig 5J’, Fig S6G). We therefore conclude that the expansion of the VA1v glomerulus during knock down of *DIP-η* is likely due to a defect in the development of glomerular morphology and positioning, rather than their maintenance.

**Figure 5:**
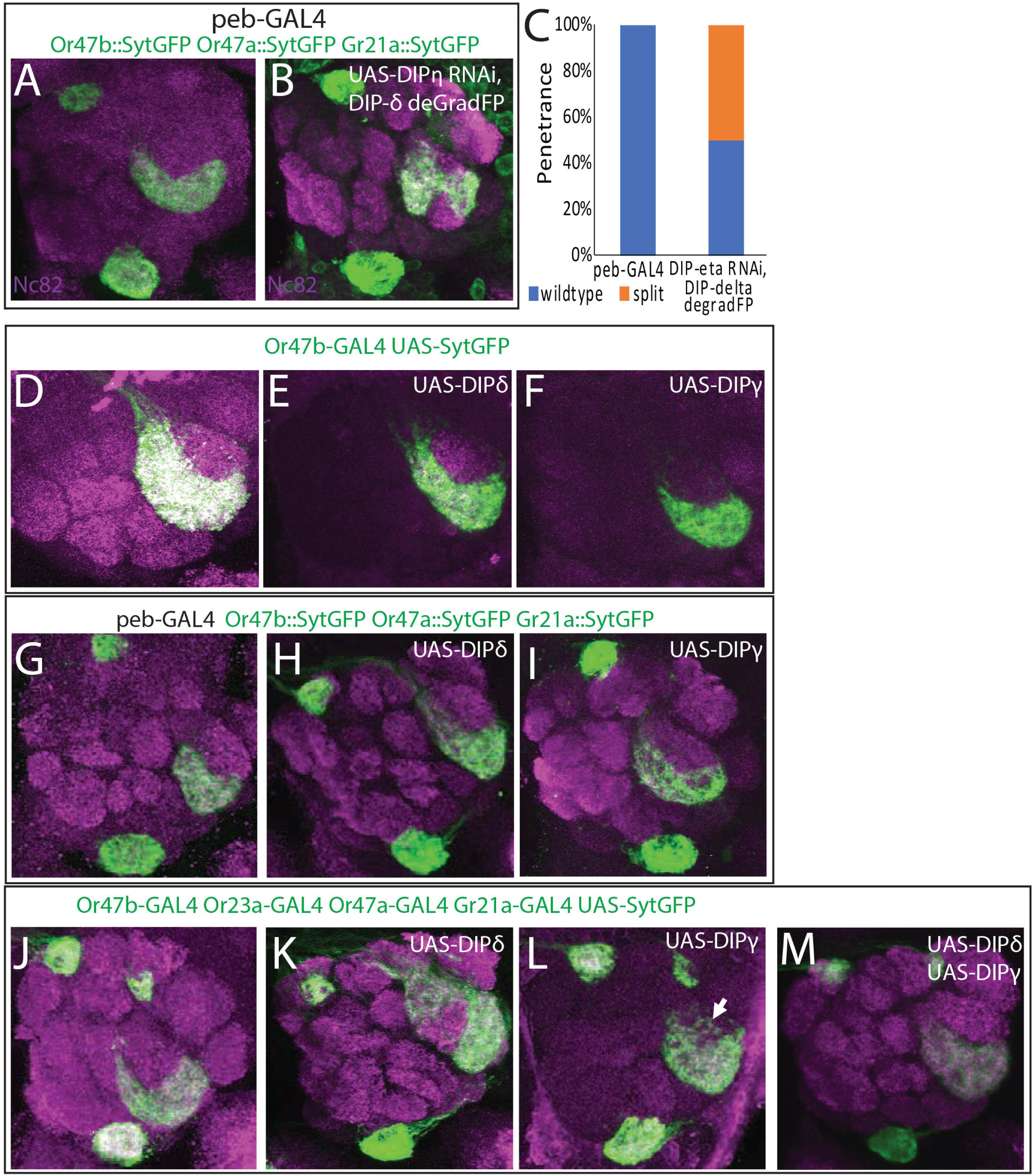
Combinatorial knock down and overexpression of *DIPs* causes cell non-autonomous defects in VA1v glomerular organization. (A, B) Knockdown of *DIP-η* and depletion of DIP-δ protein using the DeGradFP system causes the splitting of specifically the Or47b glomerulus, but not the Gr21a or Or47a glomeruli, when driven with the *peb-GAL4*. (C) Penetrance of the *DIP-η/DIP-δ* knockdown phenotype. 50% of individuals displayed a split of the Or47b glomerulus in one or both antennal lobes. (D-F) Overexpression of *DIP-δ* (E) and *DIP-γ* (F) in Or47b ORNs (green). Overexpression of either gene does not disrupt the VA1v glomerulus. (G-I) When *DIP-δ* and *DIP-γ* were over expressed in all ORNs using the *peb-GAL4* driver, no disruptions were observed in the target glomeruli for Or47b, Or47a or Gr21a ORNs (green). (J-M) *OR-GAL4*s were used to drive expression of *UAS-DIP-δ* (K), *UAS-DIP-γ* (L) and *UAS-syt-GFP* (green), and both *UAS-DIP-δ* and *UAS-DIP-γ* simultaneously (M). Mis-expression of *DIP-δ* in Or47b ORNs caused the target glomerulus to deform and split (K), while other ORNs mis-expressing *DIP-δ* were unaffected. Over expression of *DIP-γ* in Or47b ORNs caused their axons to partially invade the Or88a target glomerulus (L), like knock down of *DIP-η* (Figure 4). In contrast, other glomeruli that over expressed *DIP-γ* were unaffected. Surprisingly, overexpression of both *DIP-δ* and *DIP-γ* returned the shape of the VA1v glomerulus to its wildtype morphology (M).

We next tested whether differences in the levels of *DIP-η* would alter the severity and penetrance of the *DIP-η* knock down phenotype. We raised flies at 18 and 22 °C to reduce the level of RNAi knock down of *DIP-η*. Since RNAi expression relies on the yeast GAL4-UAS system, decreasing the temperature can generate different levels of RNAi expression and knock down of *DIP-η.* Surprisingly, flies raised at 22 °C displayed splitting of the VA1v glomerulus, in contrast with flies raised at 28 °C, which displayed an expansion of the VA1v glomerulus dorsally (n=17 65% penetrance, Fig S7I). Flies raised at 18 °C also occasionally presented split VA1v glomeruli, although at a much reduced penetrance, but mostly displayed wildtype VA1v morphology (n=14 29% penetrance, Fig S7H). These results suggest that different levels of *DIP-η* knock down is associated with qualitative differences in the position of ectopic Or47b ORN projections, which is in agreement with a model involving differential adhesion as a strategy to sort out ORN axons into distinctly positioned glomeruli.

Or47b and Or88a ORNs share all of the DIP/Dpr combinations, except for *dpr5,* which is only expressed in Or88a ORNs. We therefore hypothesized that loss of *DIP-η* in Or88a ORNs could disrupt the morphology of the VA1d glomerulus. Initially, we drove RNAi against *DIP-η* using the *Or88a-GAL4* but did not observe any significant changes to the VA1d glomerulus (Fig S7K, L). *Or88a* expression begins late in development however, and therefore knock down of *DIP-η* may begin too late in this condition to disrupt glomerular morphology. In order to knock down expression of *DIP-η* in Or88a neurons earlier in development, we used the *Bar-GAL4* to drive RNAi against *DIP-η*. *Bar* expression begins in the antennal disc and by 40 hrs APF is expressed in Or88a, Ir84a, and Ir75d ORNs, continuing into adulthood (33). In this condition, we observed deformations of the VA1d glomerulus, characterized by aberrant ventral projections of Or88a axon terminals (n=10 60% penetrance, Fig 4K, K’). These deformations were similar to the behavior of the VA1d glomerulus when the VA1v glomerulus splits. At this time, we cannot determine whether the disruption to the VA1d glomerulus is due to the knock down of *DIP-η* in multiple classes of ORNs or to the knock down beginning early in development. Nonetheless, we conclude that knock down of *DIP-η* in multiple classes of ORNs beginning early in development is sufficient to disrupt the glomerular morphology and positioning of the VA1d glomerulus.

Interestingly, *dpr5,* a ligand for *DIP-η* and *θ*, is the only gene that we assayed that is expressed in Or88a neurons but not Or47b ORNs. Knock down of *dpr5* with the *peb-GAL4* did not produce any changes to VA1v glomerular morphology (Fig S6H). This is likely due to redundancy of interactions between DIP-η/θ and Dpr2/3 or other as of yet unidentified CSRs that are also involved in controlling glomerular morphology.

### DIP/Dpr family members interact genetically and combinatorially to organize ORN axon terminals and glomerular position

Lack of wiring defects in single *DIP/dpr* knock downs might be due to the combinatorial and functionally redundant nature of DIP/Dpr family members. One approach to circumvent this problem is to knock down *DIPs/dprs* in different combinations. To investigate this, we next analyzed double knock down of *DIP-η* and *DIP-δ*. Knock down of *DIP-η* alone produces no observable defects when driven in all ORNs (Fig S6B, C). *DIP-δ*, as well as its interaction partner *dpr12* are expressed in all four glomeruli. No RNAi line currently exists for *DIP-δ*, so we instead used the deGradFP system to reduce GFP tagged DIP-δ protein in ORNs. The deGradFP construct can be driven using the GAL4-UAS system and specifically targets GFP tagged proteins for degradation by the proteasome (46). The deGradFP system utilizes a single antibody domain fragment called VhhGFP4, which binds to GFP, Venus, YFP and EYFP, fused to an E3 ubiquitin ligase. This protein, when expressed, ubiquitinates proteins tagged with GFP, targeting them for degradation (46). We combined the *peb-GAL4* mediated RNAi knock down of *DIP-η* expression with *UAS-deGradFP* expression to reduce DIP-δ protein levels in all ORNs. Analysis of Or47b, Or47a, and Gr21a axon terminals using promoter fusion constructs showed very specific and localized defects in the VA1v glomerulus (Fig 5A, B). Most noticeably, Or47b ORN axon terminals extended radially towards the DC3 and VL2a glomeruli, targeted by Or83c and Ir84a ORNs, respectively, splitting the glomerulus in two (Fig 5B). 50% of individuals displayed defects in one or both antennal lobes (n=10, Fig 5C). In this circumstance, disruption of intra-class attraction among Or47b ORN axons, and Ir84a and Or83c ORN axons due to loss of *DIP-η* and *DIP-δ*, is accompanied by the decrease in the diversity of DIP/Dpr combinations. This likely causes both classes of axons to not only reduce self-adhesion, exemplified by VA1v glomerular splits, but also might decrease diversity among ORN axons of different types. Such a split could be due to greater relative attraction of Or47b axons to axons of other classes, while retaining some self-adhesion, creating effectively two VA1v glomeruli. We therefore conclude that *DIP-η* and *DIP-δ* are required for the sorting of Or47b axons in a combinatorial, context-dependent, and non-cell autonomous fashion.

### Combinatorial and differential overexpression of DIP proteins causes local axon sorting defects

So far, we have shown that DIPs/Dprs help sort Or47b, Or88a, Or83c, and Ir84a ORN axon terminals into 4 glomerular units in combinatorial, and context-dependent fashion. Because loss of DIPs/Dprs can lead to sorting defects, we hypothesized that addition of new factors to the code would also disrupt the wiring of these glomeruli. We expected that over expression should cause ORN axons from different classes to converge because their code of expression has become more similar or interactive. To test this hypothesis, using the *Or47b-GAL4* driver we overexpressed *DIP-δ* and *DIP-γ*, a DIP which is not expressed in any of the 4 ORNs, but that interacts with Dpr11 found in all 4 (Fig 5D-F).

Overexpression of either *DIP-δ* or *DIP-γ* in Or47b neurons produced no discernable changes in the VA1v glomerulus (n=10 and n=6 respectively, Fig 5D-F). Broad overexpression of either gene with the *peb-GAL4* also produced no change in the VA1v glomerulus or in the DM3 and V glomeruli (Fig 5G-I). We reasoned that overexpression in all ORNs or in a single class might not generate enough of a contextual change in ORN-specific combinations to show a phenotype. To generate a combinatorial and a contextual overexpression, we chose to overexpress each gene in a subset of ORNs targeting distant sites in the antennal lobe. We first overexpressed *DIP-δ* with the *Gr21a, Or47a, Or23a,* and *Or47b-GAL4s* (Fig 5J-M). In this condition, we found that VA1v glomerulus became deformed and split apart (Fig 5K). We observed a large group of axons closer to the VL2a glomerulus than normal, with a smaller but connected group on the opposite side of the VA1d glomerulus near the DC3 glomerulus (Fig 5K). 80% of individuals had a disrupted VA1v glomerulus in one or both antennal lobes (n=14, Fig S7A).

Like knock down of *DIP-η*, overexpression of *DIP-γ*, in Gr21a, Or23a, Or47a and Or47b neurons, caused Or47b ORN axons to invade the VA1d glomerulus, although this invasion was partial (Fig 5L) and seen only in 50% of individuals (n=8, Fig S7A). We reasoned that because this phenotype only arose when *DIP-γ* is expressed in four classes of neurons, the *DIP-γ* now present on Or47a, Or23a, and Gr21a axons generates new interaction forces that pull Or47b axons towards the VA1d glomerulus. Or47b ORN projection defects in single overexpression of either *DIP-δ* or *DIP-γ* in Gr21a, Or23a, Or47a and Or47b neurons, suggest that cell non-autonomous effects exerted onto Or47b ORN axons by a subset of ORNs influences glomerular positioning and morphology.

We next wanted to investigate how the axon sorting forces generated by simultaneous overexpression of *DIP-δ* and *DIP-γ* interact. We therefore expressed both *DIP-δ* and *DIP-γ* in Or47a, Or23a, Gr21a, and Or47b neurons. Surprisingly, this condition rescued both phenotypes from *DIP-δ* and *DIP-γ* overexpression and restored the VA1v glomerulus to its wildtype shape (n=8, Fig 5M). This suggested to us that the differential forces generated by adding each of these factors individually were canceled out when they were overexpressed at the same time. We therefore conclude that context-dependent and combinatorial DIP/Dpr interactions generate differential forces within the antennal lobe among ORN axons to regulate glomerular formation and positioning.

### DIPs and Dprs mediate adhesive interactions between ORN axons

Our previous experiments suggested that manipulation of DIP/Dprs causes two kinds of disruption of local axon sorting: splitting of glomeruli, or expansion and invasion of a neighboring glomerulus. It is still unclear however, whether these phenotypes arise because DIPs and Dprs mediate repulsive or adhesive interactions. To investigate whether DIP-Dpr interactions are adhesive or repulsive, we expressed *dpr1* in ORNs. *dpr1* is not expressed by any class of ORNs but interacts with both DIP-η and DIP-θ, which are broadly expressed in most classes. We are therefore introducing a novel interaction into the olfactory system which allows for better study of DIP-Dpr interactions. Initially, we expressed *dpr1* using the Or47b-GAL4. We expected that if DIP-Dpr interactions were repulsive, that the VA1v glomerulus would be disrupted and that Or47b axons would diffuse beyond their glomerular boundary. In this condition, we observed no major changes to the VA1v glomerulus (n=14, Fig 6A, B). Because expression of *dpr1* in all Or47b ORNs may have been insufficient to change the context of DIP-Dpr interactions, we next expressed *dpr1* in a portion of Or47b neurons. Here we labeled all Or47b ORNs using a Or47b promoter fusion transgene driving the expression of GFP while using MARCM (Fig 5C, D). In this manner, all Or47b neurons will be labeled by GFP, but a portion will express *dpr1* and will be additionally labeled by CD2. If Dpr1-DIP-η/θ interactions are repulsive then the VA1v glomerulus should split apart or partition *dpr1* expressing Or47b axons away from wildtype neurons. We observed no changes in the VA1v glomerulus in this condition (n=5, Fig 5D). We therefore conclude that it is unlikely that DIP-Dpr interactions cause repulsion between axons.

**Figure 6:**
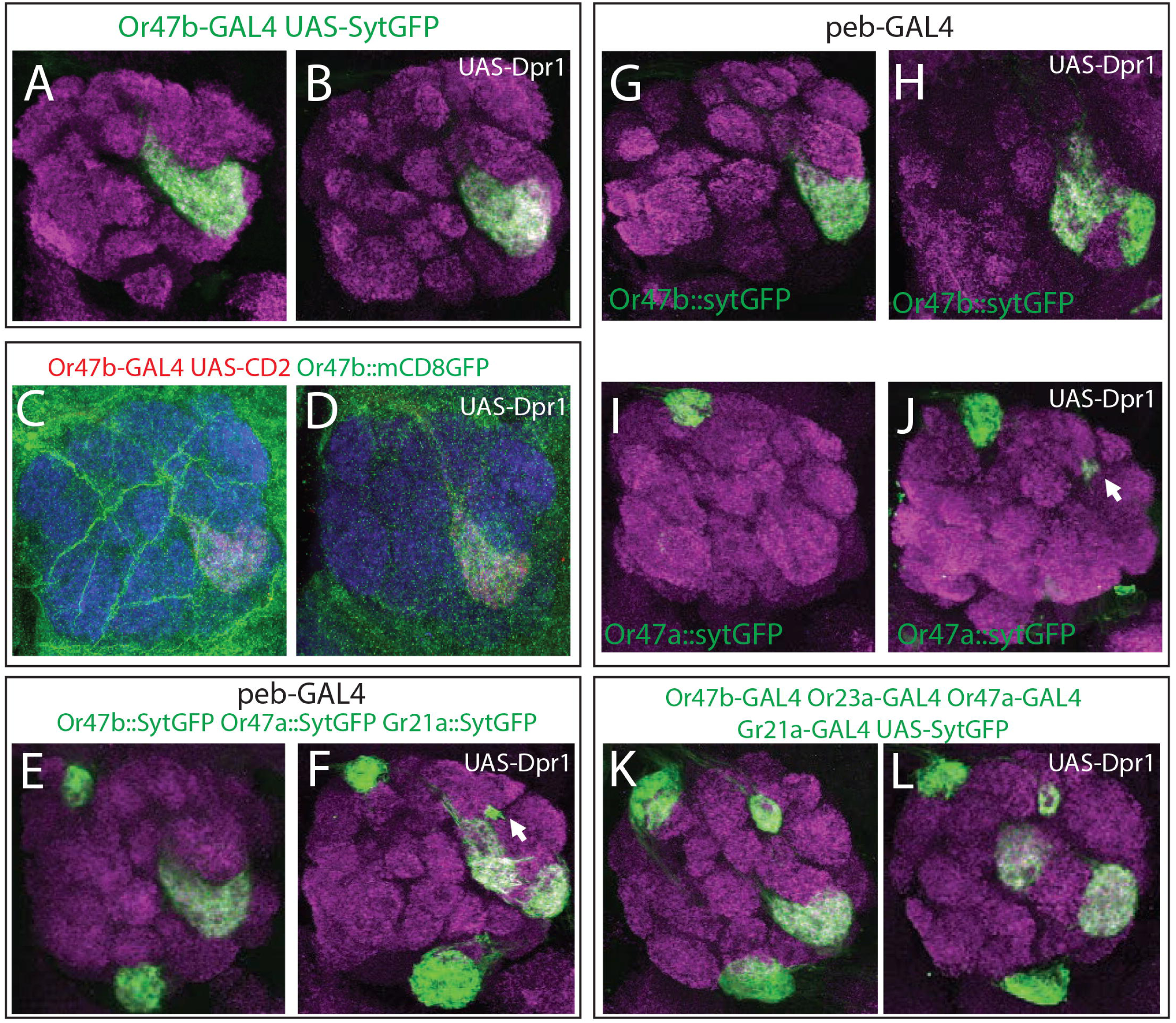
*dpr1* overexpression leads to changes in glomerular organization via adhesive interactions. (A, B) Overexpression of *dpr1* was in Or47b ORNs (green) did not perturb the VA1v glomerulus. (C, D) A subset of Or47b neurons (red) intermingle with all Or47b axons (green) in wildtype (C) and *dpr1* overexpressing flies (D). (E-J) Overexpression of *dpr1* with the *peb-GAL4* caused a split of the VA1v glomerulus (E-H). Overexpression of *dpr1* with this driver also caused mistargeting/splitting of some Or47a ORN axons (E-F, I-J), creating a highly reproducible, ectopic glomerulus. (K, L) Overexpression of *dpr1* in four classes of ORNs with *Or-GAL4* drivers caused a split Or47b glomerulus.

Our data suggest that DIPs and Dprs do not cause repulsive interactions between axons. We therefore suspected that DIPs and Dprs mediate adhesive interactions. So, we next expressed *dpr1* using the *peb-GAL4* driver to determine if DIP-Dpr interactions mediate adhesion between axons. When we visualized Or47b, Or47a, and Gr21a ORN axons, we observed that the VA1v glomerulus split (Fig 6E, F, S8B). 75% of individuals displayed a splitting of the VA1v glomerulus (n=8, Fig S8B). We also found a small and reproducible cluster of axon terminals in between the DA1 and VA1d glomeruli (Fig 6F). Labeling each ORN class individually revealed that this cluster belonged to ectopic Or47a ORN axons and was present in 50% of individuals (n=14, Fig 6G-J, S8C). These axons may mis-project along Or47a ORN axon tracks to their target DM3 glomerulus due to attraction to other ORN axons that now express *dpr1*. These data suggest that DIP-Dpr interactions likely function in ORN axon-axon adhesion.

To determine whether expression of *dpr1* in a subset of ORN classes could also disrupt the wiring of Or47a and Or47b axons we used *OR-GAL4* drivers to overexpress *dpr1* in four classes of ORNs (Or47a, Or47b, Gr21a, and Or23a, Fig 6K, L). We did not observe any change in the projection patterns of either Or47a or Or23a ORN axons. In contrast, both the V and VA1v glomeruli, targeted by Gr21a and Or47b ORNs, respectively, exhibited splitting phenotypes, stronger than ones observed in *peb-GAL4* driven overexpression (Fig 6K, S8E, F). 20% of individuals showed a split V glomerulus, and ectopic innervation of Gr21a ORN axons to a dorsal glomerulus into the antennal lobe (n=10, Fig S8D). The Gr21a ORN class is notably the only class of ORNs that does not express *DIP-η* (Fig 2H). It Is possible that these misprojected axons are attracted to dorsally neighboring axons that express *DIP-η* and therefore move dorsally towards them in the presence of Dpr1. The low penetrance and severity of the phenotype may be explained by the timing of expression of the *Gr21a-GAL4* driver. *Gr21a* expression begins relatively late in development as compared with other ORs and therefore only a few axons may express *dpr1* early enough to be affected. Splitting of the VA1v glomerulus caused some of the Or47b ORN axons to move radially and form a second glomerulus on the other side of the VA1d glomerulus (Fig 6K, L). This phenotype was observed in 40% of individuals (n=10, Fig S8B). Expression of Dpr1 in Or88a ORNs did not result in any projection defects (Fig S8G, H). These data suggest that the introduction of Dpr1 interferes with axon-axon interactions among ORNs likely through disrupting ongoing DIP/Dpr interactions.

### Dpr10 controls ORN axon guidance and axon sorting

So far, we have shown that DIPs/Dprs act primarily as adhesive molecules that regulate local ORN axon sorting. We suspected that DIPs/Dprs may regulate other aspects of ORN wiring, such as ORN-PN matching or targeting. We investigated *dpr10* because it is expressed broadly in the antennal lobe but labels the V glomerulus (Gr21a/Gr63a) most strongly. We therefore tested whether *dpr10* controls the wiring of Gr21a ORNs as well as three other classes of ORNs that are weakly labeled by *dpr10* (Or47a, Or10a, and Or47b ORNs). We used the *dpr10^MI03557^* allele which contains a MIMIC insertion, containing three premature stop codons, to analyze the loss of Dpr10 protein (38). *dpr10^MI03557^* homozygous flies die prior to eclosion, around 90hrs APF. We therefore dissected both mutant and wildtype flies at 80-90hrs APF when the antennal lobe is fully formed but prior to the death of *dpr10* mutant flies. In homozygous mutant flies, both the DM3 (Or47a) and V glomeruli were dramatically disrupted (Fig 7A-D). These defects were associated with expansion of non-converged glomeruli to neighboring glomeruli, and defects in axon guidance to ectopic sites within the antennal lobe (Fig 7A-D, S9A, B). Specifically, we observed three phenotypes for Or47a ORN axons: splitting of the DM3 glomerulus, mistargeting of a subset of Or47a ORN axons, and mistargeting to ventromedial zones in the antennal lobe (Fig 7B, S9B). Or47a ORN mistargeting phenotypes were very penetrant with >70% (n=11) of individuals showing a split or mistargeting event in one or both antennal lobes (Fig S9B). The most common phenotype observed for Gr21a ORN axons (∼39%, n=18) was a dorsal expansion of the V glomerulus (Fig 7D, S9A). In addition, we observed aberrant axons, as well as a complete splitting of the V glomerulus in a few rare cases (Fig S9A). We also analyzed Or10a axons, which are housed in the same sensillum as Gr21a neurons and target the DL1 glomerulus. Like Or47a axons, the Or10a glomerulus also exhibited splitting and dramatic mistargeting, with the most severe cases targeting to ventral positions on the opposite side of the antennal lobe, what appeared to be the VA5 glomerulus (Fig 7E, F, S9C). Some individuals displayed Or10a axons that could not form a glomerulus at all, instead, the axons formed a fascicle in between other glomeruli, mistargeting as well as losing their ability to converge (Fig 7F, S9C). These data suggest that *dpr10* not only regulates sorting of ORN axons, but also axon guidance of many ORN classes in the antennal lobe.

**Figure 7:**
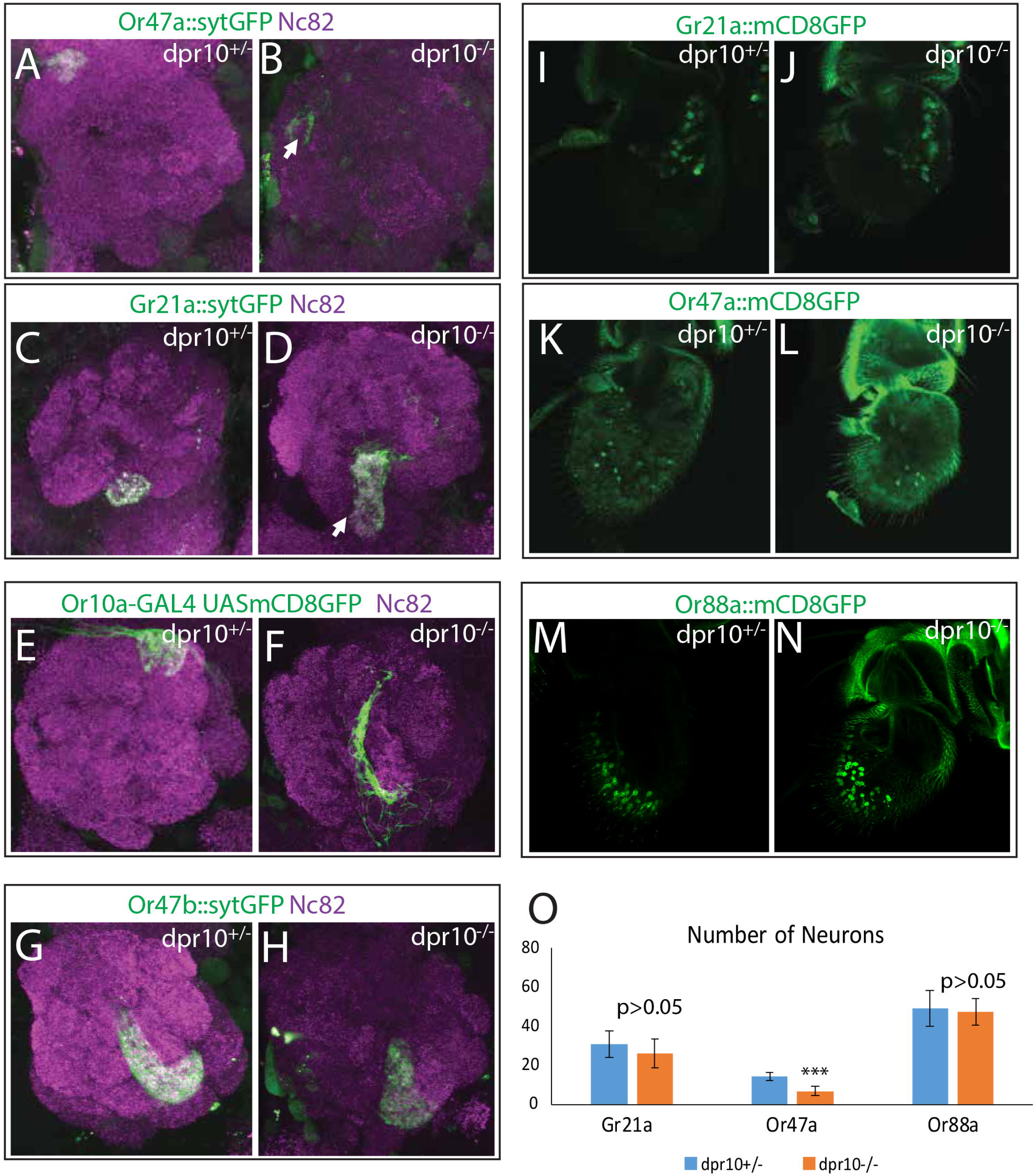
Dpr10 controls ORN wiring in the antennal lobe. Mutation of *dpr10* causes disruption of DM3 (A, B), V (C, D) and DL1 (E, F) glomeruli but not the VA1v (G, H) glomerulus as visualized with OR reporters diving *synaptotagmin-GFP* (green). Disruptions to these glomeruli included splitting of the glomerulus (B, D), mistargeting (B and F, arrow) and expansion (D, arrow). Penetrance and full summary of these phenotypes are described in Supplemental Figure 5. (G-J) ORN cell bodies were visualized in heterozygous and mutant flies for both Gr21a Or47a and Or88a ORNs using *Or-GAL4* driven *UAS-CD8GFP*. A statistically significant decrease was observed for Or47a ORNs (p<0.001, K) but not for Gr21a or Or88a ORNs (p>0.05, K).

None of the major wiring phenotypes found in Gr21a, Or47a and, Or10a ORN classes were present in the Or47b ORNs. Rather the VA1v glomerulus showed only minor disruptions of its normal shape and position (Fig 7G, H), which likely results from secondary effects of the general disorganization of the antennal lobe glomeruli in *dpr10* mutants.

Changes in the number of ORNs of a given class tends to change the size of that glomerulus (41). Thus we next wanted to confirm that the wiring phenotypes of Or47a, Or10a, and Gr21a ORNs were not due to a change in the number of ORNs in either class (33,41,44). We visualized Gr21a and Or47a ORNs in the antenna using *UAS-mCD8GFP* reporters in wildtype and *dpr10* mutants (Fig 7I-L). We observed a statistically significant reduction in the number Or47a neurons in *dpr10* mutants (14.4 vs 6.8, p<0.001, Figure 7K, L, O), suggesting that *dpr10* may regulate the survival of Or47a ORNs. This reduction may explain the reduced size of the DM3 glomerulus we observed in the antennal lobe but is insufficient to explain the misprojections of Or47a ORNs. A slight reduction of Gr21a neurons was also detected (30.8 vs 26.1) although it was not statistically significant (p>0.05, Fig 7I, J, O). Given that we observed an expansion of the Gr21a glomerulus, any reduction in the number of Gr21a neurons likely does not cause the phenotypes observed in the antennal lobe. Finally, we assayed a third class of ORNs to further investigate changes to ORN specification, Or88a (Fig 7M, N). We observed no significant change in the number of Or88a neurons in *dpr10* mutants (p>0.05, Fig 7O). These data suggest that *dpr10* mutation may partially affect the generation or survival of some classes of ORNs, in addition to its role in organizing ORN axon projections in the antennal lobe.

## Discussion

Despite our understanding of axon guidance and synaptic specificity, how complex circuits coordinate their organization across many neuronal types from a limited genetic repertoire of CSRs remains unknown. In the *Drosophila* olfactory system, 50 classes of ORNs project their axons into the antennal lobe of the brain where they connect to their partner projection neurons and organize within 50 uniquely positioned and ORN class specific glomeruli (9,11,12). Here we show that the Dpr family of Ig-domain transmembrane proteins and their heterophilic binding partners DIPs are expressed in ORN specific combinations. Mathematical analysis of class-specific DIP/Dpr expression profiles suggest ORN classes with similar DIP/Dpr profiles, can target distant glomeruli, and neighboring glomeruli are targeted by ORNs with different DIP/Dpr combinations, suggesting a role in ORN intra-class adhesion and inter-class sorting. Our results *in vivo* are in agreement with this hypothesis, as loss of a single *DIP/Dpr* gene in a specific ORN class, or multiple genes in all ORNs, causes local disruptions of ORN terminal projections and glomerular positioning, without causing defects in ORN-PN matching. Misexpression of *DIPs/dprs* causes similar phenotypes, disrupting normal axon-axon interactions in both cell autonomous and non-autonomous ways. Overexpression of DIPs/Dprs in multiple classes of ORNs can cause cell non-autonomous phenotypes in other classes of ORNs, even distant neighbors, suggesting axon-axon interactions can shift glomerular positioning. ORN projection phenotypes in different levels of single protein knock downs, and combinatorial knock down or overexpression of DIPs, suggests integration of differential adhesive forces by different DIP/Dpr combinations in ORNs contributes to glomerular structure and position. Some Dprs, Dpr10 specifically, also control additional processes during wiring, such as the fate or correct guidance of ORNs to appropriate glomerular regions, as seen in *dpr10* mutants. Together, our data reveals that DIPs/Dprs are critical players in ORN axon sorting, as well as the positioning of 50 ORN class-specific glomeruli.

Although this study provides a significant advance in understanding how axon terminals for 50 ORN classes segregate into uniquely positioned glomeruli, it is incomplete. First and foremost, we do not have expression data for several other DIP/Dpr family members (Dprs 4, 7, 14, 17, 18, 19, 20, 21, and DIPs-α and I). Second, we focused our analysis on four glomeruli targeted by Or47b, Or88a, Or83c, and Ir84a ORNs, showing only a few examples of manipulations in other ORN classes. More sophisticated systems level genetic analyses of all DIP/Dpr manipulations, in addition to identification of CSR expression profiles in each ORN class and its target projection neuron will help refine our model, in the future.

Another caveat to our analysis is that that the ORN class-specific DIP/Dpr profiles rely on MIMIC GAL4 driven reporter expression patterns in the antennal lobes. Even though these have been confirmed in the neurons of the visual system (37), it is possible that some do not reflect the endogenous gene expression. Thus, acquisition of ORN specific transcriptional profiles, together with targeted GAL4s knock-ins into individual DIP/Dpr loci will be needed in the future for a more detailed understanding of the ORN-specific DIP/Dpr combinatorial codes.

### Differential adhesion via combinatorial DIP-Dpr interactions regulate glomerular organization

One key characteristic of the peripheral olfactory system is the sorting of approximately 1500 ORN axon terminals into 50, uniquely positioned, and ORN class-specific glomerular units in the antennal lobe (9,11). Our data suggests that, in addition to guidance of ORN axons to the antennal lobe and synaptic ORN-PN matching within glomeruli, axon-axon interactions among ORNs also contribute to the glomerular organization and formation. Such axon-axon interaction among ORNs during development can function to attract and adhere ORN axon terminals of the same class, simultaneously sorting from the axon terminals of other ORN classes, which themselves must self-adhere. These interactions can also generate differential forces that also position the terminals of each ORN, and thus the position of each glomerulus with respect to others. The molecular mechanisms driving these different processes during glomerular positioning and formation are not known. The process of sorting subsets of ORN axons into different tracks successively, starts with Notch signaling and its regulation of semaphorins, as ORNs in the same sensillum are born from asymmetric divisions of the same precursor to acquire separate wiring identities (23). Each of the sibling ORNs take one of the two early axon tracks before glomeruli start to form based on their Notch state, which also determines whether or not they express Sema-2b. Sema-1a signaling also contributes to repulsion of neighboring ORN axons of different classes, but again, effects are relatively general, causing sorting defect of many ORN classes (47). In addition to repulsive signals, examples of homophilic cell adhesion proteins, such as N-Cadherin, were previously shown to regulate glomerular formation as well, by interfering with axon-axon interactions among the same class of ORNs (27,48). The effect of *Ncad* mutants however, are seen in all glomeruli, which does not explain selective adhesion that occurs among each class (27,48). Here we show that ORN-specific combinations of DIP/Dpr pairs regulate glomerular morphology and positioning within the antennal lobe. Knock down and mis-expression experiments also indicate that ORN axons interact via their DIPs/Dpr combinations suggesting that they may distinguish themselves from other ORN axons nearby, and integrate these interactions to identify and converge with ORNs of the same class, positioning the glomeruli for other ORNs with similar or slightly compatible DIP/Dpr profiles forming glomerular neighborhoods. This is particularly apparent for trichoid and coeloconic ORNs, which have more similar DIP/Dpr profiles among ORNs within each sensilla type compared to others. Given the previously reported adhesive function of DIP/Dpr interactions (35,37), our work then supports a model of differential adhesion that emerges from ORN-specific combinatorial DIP/Dpr profiles as a strategy for class-specific sorting of ORN axon terminals.

The sorting defects observed in this study could, in theory, arise due to either defects in intra-class attraction or inter-class repulsion. Taken together, our data argue for a third option, where differential adhesive interactions among axons sort out glomeruli of different classes. In many cases ligand-receptor DIP-Dpr pairs are expressed in the same ORN class, which can interact heterophilically to mediate adhesion, albeit it is also possible that some DIP/Dprs interact homophilically. Yet, some ORN classes, especially ones with more complex combinatorial codes, additionally express only specific Dpr ligands, without their DIP receptors. In these cases, we sometimes find the receptors can rather be expressed in ORN classes that target neighboring glomeruli. It is therefore possible that differential axon-axon interactions during development can help position axons from different ORN classes based on its DIP/Dpr profile. Indeed, our results with *dpr1, DIP-δ*, and *DIP-γ* overexpression supports this model of interaction. When expressed in only Or47b neurons, these genes yield no significant changes to the Or47b ORN projections or the VA1v glomerulus. Yet only when they are expressed in other ORN classes Or47b projection defects appear. This suggests that during development Dpr1, DIP-δ, and DIP-γ in Or47b neurons and other ORN classes interact with the matching DIPs or Dprs on nearby axons of other ORN classes. These interactions likely generate forces that pull out Or47b axons to distinct directions. Additional support for differential adhesion comes from the titration levels of *DIP-η* knock down, where increasing the temperature and thus the strength of knock down is accompanied by increased severity of the Or47b ORN projection phenotype. This is likely due to differing levels of self-adhesion, where a slight decrease in self adhesion can lead to glomerular splits as opposed to a full expansion of VA1v glomerulus in stronger knock downs to fully overtake the VA1d glomerulus. This is also consistent with the developmental analysis of the phenotype, which reveals that glomerular splits and positional defects are apparent at 45-50hrs APF, before axons fully expand at 55hrs APF. Previous reports have observed that the number of Or47b neurons that are positive for the GFP driven by the *Or47b-GAL4* increases over pupal development (34). This would suggest that the number of Or47b neurons that lack *DIP-η* expression at mid-pupal stages are relatively few, resulting in a modest reduction of *DIP-η* expression in the population as a whole. This reduction would increase as more neurons express the *Or47b-GAL4* resulting in full invasion.

Combinatorial knock down and overexpression of *DIPs* also support a model in which DIP/Dprs mediate differential adhesion. Overexpression of either *DIP-δ* or *DIP-γ* on their own, misdirects axon terminals indifferent directions. This is likely due to different context dependent adhesive DIP/Dpr interactions among ORN axons within the antennal lobe glomeruli in each experimental condition. Interestingly, these effects are neutralized when both DIPs are expressed simultaneously, suggesting integration of different adhesive forces exerted on the ORN axon terminals for each class.

Heterophilic adhesive interactions among DIPs/Dprs are consistent with their previously reported roles in the *Drosophila* eye in layer specific matching of photoreceptor cells with their targets in the medulla (35). Mutations in *DIPs/dprs* cause photoreceptors to overshoot their targets because they lack adhesion with their postsynaptic partners (35). In our study, we did not detect any ectopic synaptic matching of Or47b ORNs with other PNs, suggesting DIP/Dprs in ORNs might function in glomerular sorting and positioning, but not ORN-PN matching. However, these experiments are restricted to the analysis of MZ19 reporter, which labels DA1, DC3, and VA1d PNs. Thus even though it is possible that DIP/Dpr interactions might play a role in ORN-PN matching for other ORN classes, our data is in agreement with a model of differential adhesion as an efficient strategy to form and segregate 50 class specific glomeruli in the antennal lobe. At each stage of the wiring program, axons can interact with their neighbors based on their DIP/Dpr profiles where highest adhesion occurs among the axons of the same ORN class, and perhaps determine the relative glomerular position of other ORN classes nearby. Superimposed onto other earlier regulators of wiring, such as Sema-1a, Sema-2a, and N-Cadherin, glomeruli can be sorted out from one another in a repeatable fashion (27,47).

### ORN wiring programs continue after the onset of OR expression

Our results suggest that loss or addition of DIPs/Dprs generally act locally in a class-specific manner during the last stages of glomerular formation. In many instances, phenotypes can be generated by knocking down or overexpressing *DIP/dpr* genes using *OR-GAL4* drivers, after the onset of olfactory receptor expression. This result is rather interesting given the current view of ORN circuit assembly. For many years, the consensus has been olfactory receptor genes are turned on after the glomerular formation is complete. Our results suggest that, at least for olfactory receptors that are turned on early in development, glomerular patterns are not entirely established by the onset of OR expression. The finding that class specific knock down and over expression experiments using *Or47b-GAL4* drivers for many *DIP* genes leads to dramatic defects in the target VA1v glomerulus, indicates ongoing axonal decisions that require DIPs/Dprs after the onset of *Or47b* expression.

In addition, developmental analysis of knock down of *DIP-η* in Or47b ORNs shows that glomerular deformation can be detected as early 45 hrs APF during the finalization of glomerular formation. This suggests that DIPs and Dprs play some role in the development of glomerular morphology and positioning, while others may be more critical for the maintenance of proper glomerular shape. This conclusion is bolstered by the observation that some DIPs and Dprs have developmentally dynamic expression patterns, with distinct expression patterns at 40-50 hrs APF, while others show little to no expression at this developmental stage. It is likely that different DIPs and Dprs play different roles in ORN wiring depending on their expression pattern and timing.

It should also be noted that some phenotypes arise when other *Or-GAL4s,* that turn on later, are used in conjunction with the *Or47b-GAL4*. This suggests that while glomeruli obtain they final shape by 48hrs APF, this morphology is not set and can be altered later in development by the mis-expression or knock down of CSRs.

When a specific *DIP/dpr* gene, or gene combination, is lost or ectopically expressed in ORNs, axons invade the glomerulus targeted by a class of ORNs with the most compatible DIP/Dpr code, and/or which also now has new adhesive properties due to the perturbed genetic state. This role is akin to Dscam, a fellow Ig superfamily protein, involved in controlling dendritic self-avoidance, where combinatorial expression of thousands of Dscam splice isoforms regulate recognition of self vs non-self-dendritic processes to produce distinct dendritic zones for each neuron (36). In the case of DIPs/Dprs in the olfactory system, combinatorial expression of DIP/Dprs regulates sorting of ORN axon terminals that belong to the same class (self) from axon terminals of other ORN classes (non-self) which target neighboring glomeruli in the antennal lobe.

### Regulation of DIP/Dpr Expression

How do the class specific patterns of DIP/Dpr expression arise? It seems likely that similar mechanisms that lead to the singular expression of a particular olfactory receptor in each neuron would also control the expression of DIPs/Dprs. There are three major modes of regulation of OR selection: 1-prepatterning of the antennal disc by transcription factor networks that determine sensilla precursor identity, 2-regulation of neuronal fates by Notch-Delta signaling during asymmetric precursor cell divisions, and 3-terminal selector transcription factors regulating olfactory receptor gene expression and possibly other ORN identifiers (19,33,49). Each mode of regulation clearly controls ORN wiring as prepatterning network mutants change ORN connectivity to converted fates (33,41), Notch pathway mutants behave similarly (19,43), and factors like Pdm3 regulate glomerular shape as well OR expression (50). It is likely that each mode of regulation, layered on top of each other, work in concert to control DIP/Dpr expression. This would lead to class-specific differences in their DIP/Dpr profiles, which would be generated by the developmental programs mediating terminal differentiation of each ORN class, increasing the complexity of the DIP/Dpr code for each ORN class. There might be some DIP/Dprs, like *DIP-η* and *dpr12*, that are expressed earlier and more abundantly in future ORNs, perhaps starting at precursor stages to coarsely sort early axons. Some DIPs/Dprs are indeed expressed in our RNA-seq analysis at 8 hrs APF although at much lower levels when compared to 40 hrs APF and adult antennae. DIP/Dprs expressed in only few classes of ORNs, might be superimposed onto existing DIP/Dpr profiles in later stages of development, as individual ORN identities are defined by the onset of OR expression. Deeper understanding of exactly which transcription factors control DIP/Dpr expression and how they relate to larger programs of neuronal specification is needed.

### Convergent molecular logic for glomerular organization

The mammalian olfactory bulb is organizationally very similar to the *Drosophila* antennal lobe. In both organisms, the ORNs that express the same receptor converge their axons onto a single class-specific glomerulus (9,51). Mammalian olfactory receptors are G-protein coupled receptors and differentially regulate the expression of adhesive and repulsive CSRs to position and sort ORN specific glomeruli using both ligand dependent and independent signaling (16,17). Differential, ligand-independent cAMP signaling from each mammalian olfactory receptor gene regulates graded expression of Semaphorin and Neuropilin in different ORN classes, and control the positioning of each glomerulus (16,17). In addition, olfactory receptor neuron activity refines glomerular convergence through differential expression of homophilic adhesion Ig-domain proteins Kirrel2/3, and repulsive transmembrane signaling proteins EphA and EphrinA, which regulate intra-class attraction and inter-class repulsion respectively (16,17).

In contrast to mammalian olfactory receptors, *Drosophila* olfactory receptors are ligand gated cation channels, and do not contribute to ORN wiring (18). Interestingly, DIPs and Dprs share homology with Kirrels and seem to operate with a similar logic. Differential and graded expression of adhesion proteins, Kirrels in mammals and a combination of DIPs/Dprs in flies, mediates class-specific glomerular convergence. They do this by regulating adhesion among the axons from the same ORN class, by creating differential adhesion forces locally in the olfactory bulb based on ORN-specific cell surface receptor profiles. Thus, even though olfactory receptors in mammals and *Drosophila* are functionally and structurally diverse, our point to a possible evolutionarily convergent downstream molecular strategies that sort ORN axon terminals into distinct glomeruli in a class specific manner.

## Materials and Methods

### Fly Genetics

OR-CD8 GFP, OR-Syt GFP, OR-GAL4, IR-GAL4, GR-GAL4 lines were from Leslie Vosshall, Barry Dickson, Richard Benton and John Carlson, respectively. Dpr-GAL4 and DIP-GAL4 lines were from Larry Zipursky. UAS-CD8 GFP, UAS>STOP>GFP, UAS-SytGFP, ey-FLP, MZ19-CD8 GFP, UAS-RFP, UAS-dpr1, UAS-DIPγ, UAS-DIPδ, dpr10^MI03557^ and UAS-RNAi lines were all from Bloomington Stock Center.

#### Fly Genotypes

Figure 1A, C, D, E. *w*^*1118*^

Figure 2A. dpr-GAL4/UAS-sytGFP

Figure 2B and S2B. eyFLP/+; UAS>STOP>GFP /+; dpr5-GAL4/+

Figure 2C and S2F. eyFLP/+; UAS>STOP>GFP /+; dpr10-GAL4/+

Figure 2D and S2G. eyFLP/+; UAS>STOP>GFP/+; dpr11-GAL4/+

Figure 2E and S2J. DIP-β-GAL4/eyFLP;; UAS>STOP>GFP/+

Figure 2F and S2K. eyFLP/+; UAS>STOP>GFP/+; DIP-γ-GAL4/+

Figure 2G and S2M. eyFLP/+; DIP-ε-GAL4/UAS>STOP>GFP

Figure 2H and S2O. eyFLP/+; DIP-η-GAL4/UAS>STOP>GFP

Figure S1A. eyFLP/+; dpr2-GAL4 UAS-CD8GFP/CyO; FRT82 GAL80/FRT82

Figure S1B and S2A. eyFLP/+; dpr3-GAL4/UAS>STOP>GFP

Figure S1C and S2C. eyFLP/+; UAS>STOP>GFP/+; dpr6-GAL4/+

Figure S1D and S2D. dpr8-GAL4/eyFLP;; UAS>STOP>GFP/+

Figure S1E and S2E. eyFLP/+; UAS>STOP>GFP/+; dpr9-GAL4/+

Figure S1F. eyFLP/+; dpr12-GAL4/UAS-sytGFP; FRT82 GAL80/FRT82

Figure S1G. dpr13-GAL4/UAS-sytGFP

Figure S1H and S2H. eyFLP/+; UAS>STOP>GFP/+; dpr15-GAL4/+

Figure S1I and S2I. eyFLP/+; UAS>STOP>GFP/+; dpr16-GAL4/+

Figure S1J and S2L. eyFLP/+; UAS>STOP>GFP/+; *DIP-δ*-GAL4/+

Figure S1K and S2N. eyFLP;/+ DIP-ζ-GAL4/UAS>STOP>GFP

Figure S1L and S2P. eyFLP/+; DIP-θ-GAL4/UAS>STOP>GFP

Figure S1M. dpr2-GAL4/UAS-CD8GFP

Figure S1N. dpr3-GAL4/UAS-CD8GFP

Figure S1O. UAS-CD8GFP/+; dpr5-GAL4/+

Figure S1P. UAS-CD8GFP/+; dpr10-GAL4/+

Figure S1Q. UAS-CD8GFP/+; dpr11-GAL4/+

Figure S1R. dpr12-GAL4/UAS-CD8GFP

Figure S1S. DIP-β-GAL4/+; UAS-CD8GFP/+

Figure S1T. UAS-CD8GFP/+; *DIP-γ*-GAL4/+

Figure S1U. UAS-CD8GFP/+; *DIP-δ*-GAL4/+

Figure S1V. DIP-ε-GAL4/UAS-CD8GFP

Figure S1W. DIP-η-GAL4/UAS-CD8GFP

Figure S3A. dpr-GAL4/UAS-DenMark

Figure S3B. dpr2-GAL4/UAS-DenMark

Figure S3C. dpr3-GAL4/UAS-DenMark

Figure S3D. UAS-DenMark/+; dpr6-GAL4/+

Figure S3E. UAS-DenMark/+; dpr10-GAL4/+

Figure S3F. dpr12-GAL4/UAS-DenMark

Figure S3G. dpr8-GAL4/+; UAS-DenMark/+

Figure S3H. DIP-α-GAL4/+; UAS-DenMark/+

Figure S3I. UAS-DenMark/+; *DIP-γ*-GAL4/+

Figure S3J. DIP-η-GAL4/UAS-DenMark

Figure S3K. dpr13-GAL4/UAS-DenMark

Figure S3L. DIP-ζ-GAL4/UAS-DenMark

Figure S6A. peb-GAL4/+; Or47b-sytGFP, Or47a-sytGFP, Gr21a-sytGFP/+

Figure S6B. peb-GAL4/+; Or47b-sytGFP, Or47a-sytGFP, Gr21a-sytGFP/UAS-DIP-α RNAi

Figure S6C. peb-GAL4/+; Or47b-sytGFP, Or47a-sytGFP, Gr21a-sytGFP/+; UAS-dpr12 RNAi/+

Figure S6D. peb-GAL4/+; Or47b-sytGFP, Or47a-sytGFP, Gr21a-sytGFP/UAS-DIP-η RNAi

Figure S6E. peb-GAL4/+; Or47b-sytGFP, Or47a-sytGFP, Gr21a-sytGFP/+; UAS-dpr8 RNAi/+

Figure S6F. peb-GAL4/+; Or47b-sytGFP, Or47a-sytGFP, Gr21a-sytGFP/+; UAS-DIP-θ RNAi/+

Figure S6G. peb-GAL4/+; Or47b-sytGFP, Or47a-sytGFP, Gr21a-sytGFP/+; UAS-dpr20 RNAi/+

Figure S6H. peb-GAL4/+; Or47b-sytGFP, Or47a-sytGFP, Gr21a-sytGFP/+; UAS-dpr5 RNAi/+

Figure S6I. peb-GAL4/+; Or47b-sytGFP, Or47a-sytGFP, Gr21a-sytGFP/+; UAS-dpr9 RNAi/+

Figure S6J. peb-GAL4/+; Or47b-sytGFP, Or47a-sytGFP, Gr21a-sytGFP/+; UAS-dpr10 RNAi/+

Figure 4A. Or47b-GAL4 UAS-sytGFP/CyO

Figure 4B. Or47b-GAL4 UAS-sytGFP/UAS-DIP-η RNAi

Figure 4C-D. Or47b-GAL4 UAS-RFP/CyO; Or88a-mCD8GFP/+

Figure 4C-E’. Or47b-GAL4 UAS-RFP/UAS-DIP-η RNAi; Or88a-mCD8GFP/+ Figure 4F. Or47b-GAL4 UAS-RFP MZ19-mCD8GFP/CyO

Figure 4F’. Or47b-GAL4 UAS-RFP MZ19-mCD8GFP/UAS-DIP-η RNAi Figure 4H-J. Or47b-GAL4 UAS-RFP/CyO

Figure 4H’-J’. Or47b-GAL4 UAS-RFP/UAS-DIP-η RNAi

Figure 4K. Bar-GAL4/+; UAS-mCD8GFP/+

Figure 4K’. Bar-GAL4/+; UAS-mCD8GFP/UAS-DIP-η RNAi

Figure S7A. Or47b-GAL4 Or47a-GAL4 Or23a-GAL4 Gr21a-GAL4 UAS-sytGFP/CyO; TM2/TM6B

Figure S7B. Or47b-GAL4 Or47a-GAL4 Or23a-GAL4 Gr21a-GAL4 UAS-sytGFP/ UAS-DIP-η RNAi; TM2/TM6B

Figure S7D. Or47b-GAL4 UAS-RFP/CyO; Or88a-mCD8GFP/+

Figure S7E. Or47b-GAL4 UAS-RFP/UAS-DIP-η RNAi; Or88a-mCD8GFP/+

Figure S7H and I. Or47b-GAL4 UAS-RFP/UAS-DIP-η RNAi; Or88a-mCD8GFP/+

Figure S7K. Or88a-GAL4 UAS-CD8GFP/CyO

Figure S7L. Or88a-GAL4 UAS-CD8GFP/UAS-DIP-η RNAi

Figure 5A. peb-GAL4/+; Or47b-sytGFP, Or47a-sytGFP, Gr21a-sytGFP/+

Figure 5B. peb-GAL4/+; Or47b-sytGFP, Or47a-sytGFP, Gr21a-sytGFP/UAS-DIP-η RNAi; DIP-δ-GFP/UAS-deGradFP

Figure 5D. Or47b-GAL4 UAS-SytGFP/CyO; TM2/TM6B

Figure 5F. Or47b-GAL4 UAS-SytGFP/+; UAS-DIP-δ/TM6B

Figure 5E. Or47b-GAL4 UAS-SytGFP/+; UAS-DIP-γ/TM6B

Figure 5G. peb-GAL4/+; Or47b-sytGFP, Or47a-sytGFP, Gr21a-sytGFP/+

Figure 5H. peb-GAL4/+; Or47b-sytGFP, Or47a-sytGFP, Gr21a-sytGFP/+; UAS-DIP-δ/+

Figure 5I. peb-GAL4/+; Or47b-sytGFP, Or47a-sytGFP, Gr21a-sytGFP/+; UAS-DIP-γ/+

Figure 5J. Or47b-GAL4 Or47a-GAL4 Or23a-GAL4 Gr21a-GAL4 UAS-sytGFP/CyO; TM2/TM6B

Figure 5K. Or47b-GAL4 Or47a-GAL4 Or23a-GAL4 Gr21a-GAL4 UAS-sytGFP/+; UAS-DIP-δ/TM6B

Figure 5L. Or47b-GAL4 Or47a-GAL4 Or23a-GAL4 Gr21a-GAL4 UAS-sytGFP/+; UAS-DIP-γ/TM6B

Figure 5M. Or47b-GAL4 Or47a-GAL4 Or23a-GAL4 Gr21a-GAL4 UAS-sytGFP/+; UAS-DIP-γ/UAS-DIP-δ

Figure 6A. Or47b-GAL4 UAS-sytGFP/CyO

Figure 6B. Or47b-GAL4 UAS-sytGFP/UAS-Dpr1

Figure 6C. eyFLP/+; Or47b-GAL4 UAS-CD2 Or47b-mCD8GFP/CyO; FRT82 GAL80/FRT82

Figure 6D. eyFLP/+; Or47b-GAL4 UAS-CD2 Or47b-mCD8GFP/UAS-dpr1; FRT82 GAL80/FRT82

Figure 6E. peb-GAL4/+; Or47b-sytGFP, Or47a-sytGFP, Gr21a-sytGFP/CyO

Figure 6F. peb-GAL4/+; Or47b-sytGFP, Or47a-sytGFP, Gr21a-sytGFP/UAS-dpr1

Figure 6E. peb-GAL4/+; Or47b-sytGFP/CyO

Figure 6F. peb-GAL4/+; Or47b-sytGFP/UAS-dpr1 Figure 6E. peb-GAL4/+; Or47a-sytGFP/CyO

Figure 6F. peb-GAL4/+; Or47a-sytGFP/UAS-dpr1

Figure 6K and S7E. Or47b-GAL4 Or47a-GAL4 Or23a-GAL4 Gr21a-GAL4 UAS-sytGFP/CyO

Figure 6L and S7F. Or47b-GAL4 Or47a-GAL4 Or23a-GAL4 Gr21a-GAL4 UAS-sytGFP/UAS-Dpr1

Figure S8G. Or88a-GAL4 UAS-CD8GFP

Figure S8H. Or88a-GAL4 UASCD8GFP/UAS-Dpr1

Figure 7A. Or47a-sytGFP; *dpr10^MI03557^*/TM6B

Figure 7B. Or47a-sytGFP; *dpr10^MI03557^*

Figure 7C. Gr21a-GAL4 UAS-sytGFP; *dpr10^MI03557^*/TM6B

Figure 7D. Gr21a-GAL4 UAS-sytGFP; *dpr10^MI03557^*

Figure 7E. Or10a-GAL4 UAS-CD8GFP; *dpr10^MI03557^*/TM6B

Figure 7F. Or10a-GAL4 UAS-CD8GFP; *dpr10^MI03557^*

Figure 7G. Or47b-sytGFP; *dpr10^MI03557^*/TM6B

Figure 7H. Or47b-sytGFP; *dpr10^MI03557^*

Figure 7I. Or47a-GAL4 UAS-sytGFP/UASCD8GFP; *dpr10^MI03557^*/TM6B

Figure 7J. Or47a-GAL4 UAS-sytGFP/UASCD8GFP; *dpr10^MI03557^*

Figure 7K. Gr21a-GAL4 UAS-CD8GFP; *dpr10^MI03557^*/TM6B

Figure 7L. Gr21a-GAL4 UAS-CD8GFP; *dpr10^MI03557^*

Figure 7M. Or88a-GAL4 UAS-CD8GFP; *dpr10^MI03557^*/TM6B

Figure 7N. Or88a-GAL4 UAS-CD8GFP; *dpr10^MI03557^*

### RNA-seq

RNAseq was performed as described before. Wandering third instar larval antennal discs (∼70 for each genotype), 8hr APF pupal antennae (∼50 for each genotype), 40hr APF pupal antennae (∼50 for each genotype), and adult antennae (150 males and 150 females) from w1118 flies were dissected. We extracted RNA only from the antennal portion of the larval eye-antennal discs in order to remove contamination by transcripts from the developing eye. RNA sequencing libraries were prepared with TruSeq Stranded mRNA Sample Prep Kit (Illumina) following the manufacturer’s instructions. For the RNA fragmentation step, 94°C, 2min was used with the intention to obtain a median size ∼185bp. PCR amplification was done with 15 cycles. A total of 24 multiplexed libraries (barcoded) were accessed for quality and mixed altogether before separating to two identical pooled libraries, which are subject to cluster generation followed by Illumina 50bp paired-end sequencing by UNC High-Throughput Sequencing Facility (HTSF), as described in (33).

### Analysis of RNAseq data

Following Li et al., The *Drosophila melanogaster* transcriptome (r5.57) was downloaded from Flybase.org and an indexed was created with bwa-0.7.8 (52). Each sequencing file was aligned to the transcriptome, and .sam files for each sample were generated. At least 80% of the total reads were able to align to the reference sequence. Count tables were then made for each sample using featureCount and a customized python script, and further consolidated into a matrix containing transcript ID and read counts from all genotypes for each stage with a Ruby script (53). These matrices were used as inputs for differential expression analysis using customized DESeq2 R script (54).

### Immunohistochemistry

Samples were fixed with 4% paraformaldehyde, washed with phosphate buffer with 0.2% Triton X-100, and staining as previously described. Primary antibodies were used in the following dilutions: rabbit α-GFP 1:1000 (Invitrogen), rat α-Ncad 1:20 (Developmental Studies Hybridoma Bank), mouse α-Bruchpilot 1:50 (Developmental Studies Hybridoma Bank), mouse α-rat CD2 1:200 (Serotec), rabbit α-RFP 1:200 (), chicken α-GFP 1:700 (). The following secondary antibodies were used: Alexa 488 goat α-rabbit 1:1000, goat α-mouse-Cy3 1:100, Alexa 568 goat α-mouse IgG highly cross-adsorbed 1:300, Alexa 647 goat α-rat 1:200, Alexa 633 goat α-mouse 1:200, goat α-rabbit Cy3 1:200, Alexa 488 goat α-rat 1:200, Alexa 488 goat α-chicken 1:700, goat α-rat Cy3 1:200. Confocal images were taken by an Olympus Fluoview FV1000 or Zeiss LSM 510 (Light Microscopy Core Facility).

### Statistical analysis of ORN class specific DIP/Dpr profiles

Biclustering analysis was performed using existing “biclust” package in R to hierarchically cluster both DIP/Dpr expression profile in each ORN class, and the ORNs that express each DIP or Dpr gene. Multidimensional scaling (MDS) was performed in R on a matrix for each ORN class and their DIP/Dpr profiles in a binary fashion (0, 1), where value 1 represents presence of a given DIP or Dpr in a given ORN class. The results were plotted in two dimensions.

k-means clustering in R was used to identify the 10 clusters below, which were later used to color code each cluster on the MDS plot. Same color coding was used on the antennal lobe scheme to highlight actual glomerular positions for each ORN class.

Clusters are:

1- Ir75abc2, Or85d, Or49a85f, Or67a, Or10a,

2- Ir92a76a, Or35a, Ir84a, Ir76ab, Or67b, Or67c, Gr21a, Or2a,

3- Or42a, Or49b, Or85b, Or7a, Or47a33b, Or83c, Or43a,

4- UNKNOWN1, Or42b, Or85a,

5- Ir75d, Or43b, Or56a33a, Or22ab, Or67d,

6- Ir64a, Ir75abc, Or46a, Ir31a, Ir75a, Ir41a, Or98a,

7- UNKNOWN2, Or59c, Or71a, Or82a, Or9a, Or59b, Or23a,

8- Or33c, Or13a, Or69ab, Or92a, Or47b, Or65abc, Or88a, Or19a

## Acknowledgements

We would like to thank Hugo Bellen, Kai Zinn and Larry Zipursky for generously sharing reagents and input into this study. We thank Bloomington Stock Center and Drosophila Genetic Resource Center for their services. We acknowledge the services provided by Duke Light Microscopy Core Facility, Duke Microarray Core Facility, UNC High-Throughput Sequencing Facility.

**Author Contributions**
SB and PCV conceptualized the experiments and wrote the manuscript. SB preformed the experiments, with the help from SN, IS, and SB. SB and PCV analyzed the data, with help from SN and IS. Bioinformatics and mathematical analyses were performed by SB and PCV with the help of CDJ and SM, respectively.

